# A Newborn F-box Gene Blocks Gene Flow by Selectively Degrading Phosphoglucomutase in Species Hybrids

**DOI:** 10.1101/2024.09.11.612556

**Authors:** Dongying Xie, Yiming Ma, Pohao Ye, Yiqing Liu, Qiutao Ding, Gefei Huang, Marie-Anne Felix, Zongwei Cai, Zhongying Zhao

**Affiliations:** Department of Biology, Hong Kong Baptist University, Hong Kong SAR; Department of Chemistry, Hong Kong Baptist University, Hong Kong SAR; Institut de Biologie de l’Ecole Normale Supérieure, CNRS, Inserm Paris France; State Key Laboratory of Environmental and Biological Analysis, Hong Kong Baptist University, Hong Kong SAR

**Keywords:** Gene duplication, Bateson-Dobzhansky-Muller incompatibility, F-box gene, *Caenorhabditis* nematodes, Species hybrid

## Abstract

The establishment of reproductive barriers such as postzygotic <u>h</u>ybrid incompatibility (HI) remains the key to speciation. Gene duplication followed by differential functionalization has long been proposed as a major model underlying HI, but few supporting evidence exists. Here, we demonstrate that a new-born F-box gene, *Cni-neib-1*, of the nematode *Caenorhabditis nigoni* specifically inactivates an essential phosphoglucomutase encoded by *Cbr-shls-1* in its sister species *C. briggsae* and their hybrids. Zygotic expression of *Cni-neib-1* specifically depletes *Cbr-*SHLS-1, but not *Cni-*SHLS-1, in approximately 40 minutes starting from gastrulation, causing embryonic death. *Cni-neib-1* is one of thirty-three paralogues emerging from a recent surge in F-box gene duplication events within *C. nigoni*, all of which are evolving under positive selection. *Cni-neib-1* undergoes turnover even among *C. nigoni* populations. Differential expansion of F-box genes between the two species could reflect their distinctive innate immune responses. Collectively, we demonstrate how recent duplication of genes involved in protein degradation can cause unintended destruction of targets in hybrids that leads to HI, providing an invaluable insight into mechanisms of speciation.

**Significance statement:** Understanding how reproductive isolation forms and triggers speciation remains a central topic in evolutionary biology. In this study, the authors reveal a novel molecular mechanism that obstructs gene flow between the nematodes *Caenorhabditis briggsae* and *C. nigoni*. A recently evolved, species-specific F-box gene in *C. nigoni* selectively depletes an essential phosphoglucomutase enzyme in *C. briggsae*, leading to embryonic lethality in hybrids. This mechanism effectively prevents gene flow from *C. nigoni* to *C. briggsae*. This discovery underscores how a newborn gene, arising from recent gene duplication, can contribute to reproductive isolation by inadvertently disrupting a functionally important protein in hybrid offspring.

## Introduction

The formation of reproductive barriers such as postzygotic hybrid incompatibility (HI) is a fundamental process in speciation because it prevents gene flow between individuals (1). HI is usually caused by genes that undergo rapid evolution (2). Such gene can arise from gene duplication events which frequently lead to rapid divergence in gene functions while generating evolutionary novelties (3–5). For example, gene duplication followed by translocation and inactivation of different copies in two lineages can lead to inviable hybrids inheriting both non-functional copies (6–8). In theory, HI may derive not only from differential inactivation of functional gene duplicates, but also through neofunctionalization of duplicated genes (3). This can occur, for instance, through a negative epistatic interaction in the hybrids between a newly duplicated allele from one parental species and an existing allele from the other. However, mapping and cloning of alleles involved in HI is a challenging task (4, 5). Therefore, despite the prevalence of recent and differential gene duplications even between closely related species (9), no empirical evidence supports this prediction.

The free-living nematodes, *Caenorhabditis briggsae* and *C. nigoni*, both closely related to the model organism *C. elegans*, represent an ideal model for investigating HI mechanisms, especially whether gene duplication shapes reproductive isolation. First, extensive gene duplications are frequently observed across *Caenorhabditis* species, resulting in large variations in their genome sizes (10). This is particularly evident between *C. briggsae* and *C. nigoni*, with the latter having a genome and proteome 22% and 35% larger, respectively, than those of the former (11, 12) (Fig. 1A). Despite these differences, the two species can still mate and produce partially viable progeny (13, 14). Second, parallelling the asymmetry in genome sizes and gene numbers, asymmetric gene flow has been observed between the two species: gene flow from *C. briggsae* to *C. nigoni* is permitted, while reciprocal gene flow is blocked (13, 15, 16). Specifically, backcrossing of their F1 hybrids to *C. nigoni* produces viable F2 progeny (denoted as nB2 hybrids), whereas backcrossing to *C. briggsae* results in no viable progeny, with all the hybrids progeny (denoted as bB2 hybrids) being arrested as embryos (Fig. 1A). Thus, hybrid viability is expected to strongly rely on certain *C. nigoni* genomic regions. Third, a markedly increased gene dysregulation is observed in bB2 hybrid embryos compared to F1 or nB2 hybrid embryos (15), suggesting the manifestation of potential negative epistatic interactions in the regulation of conserved gene expression.

**Figure 1.**
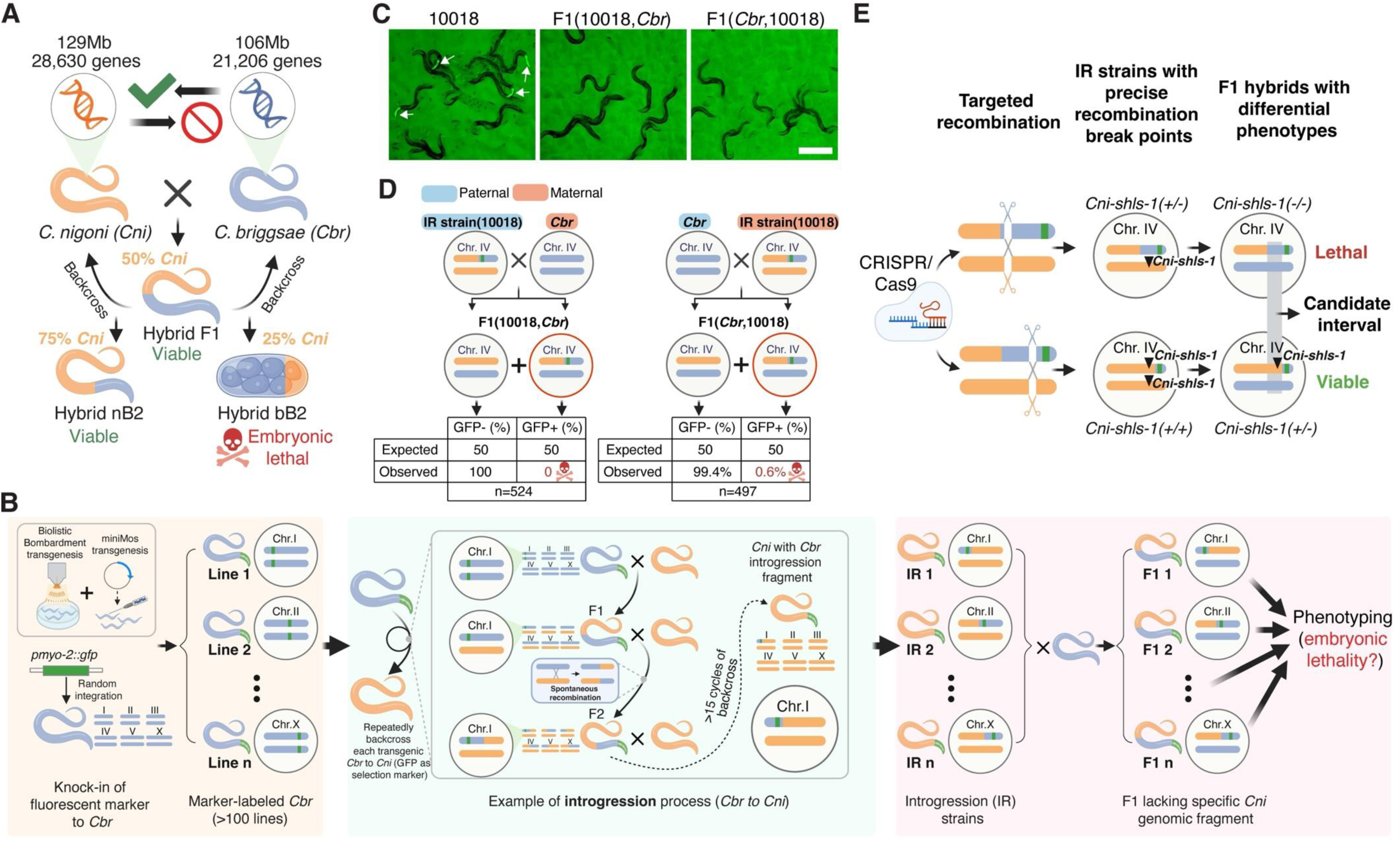
Absence of a hybrid lethality suppressor locus on *C. nigoni* Chr. IV blocks gene flow from *C. nigoni* to *C. briggsae*. **(A)** Schematic illustrating the asymmetrical gene flow between *C. nigoni* (*Cni*, orange) and *C. briggsae* (*Cbr,* blue). Viable progeny result from backcrossing F1 hybrids with *C. nigoni,* but not with *C. briggsae*. The backcrossed hybrids to *C. nigoni* and *C. briggsae* are denoted as nB2 and bB2, respectively. **(B)** Schematic showing the methodology to identify the hybrid lethality suppressor loci by phenotyping F1 hybrids lacking *C. nigoni* genomic fragment. Over 100 *C. briggsae* strains with chromosomally integrated GFP markers were generated and repeatedly backcrossed with *C. nigoni*, using GFP as selection marker, to produce *C. nigoni* strains with introgressed *C. briggsae* genomic fragments (IR strains). Each IR strain was crossed with *C. briggsae*, and hybrid progeny with homozygous *C. briggsae* introgressions (lacking syntenic *C. nigoni* fragments) were phenotyped for embryonic lethality. **(C)** Representative images of adult worms from an IR strain carrying the right arm of *C. briggsae* Chr. IV, i.e., ZZY10018 (left), and its F1 hybrids with *C. briggsae* mothers (middle) or fathers (right) on culture plates. GFP expression in the pharynx is indicated by arrowheads. Scale bar: 0.5mm. **(D)** Homozygosity of the *C. briggsae* Chr. IV right arm causes complete lethality in the F1 hybrids. Shown are the expected and observed proportions of hybrid F1 adults carrying (GFP+) or lacking GFP (GFP-) marker from crosses between *C. briggsae* and ZZY10018. Genotype schematics (Chr. IV only) for parents and F1 hybrids from reciprocal crosses are shown. Note that a complete or nearly complete absence of GFP+ worms is observed when *C. briggsae* is the maternal (left) or paternal (right) parent, respectively. In all panels, F1 hybrid progeny are named by crossing direction, with paternal and maternal parents on the left and right, respectively; Strain prefixes ZZY are omitted for brevity. **(E)** Strategy for mapping the *Cni-shls-1* locus using IR strains with defined introgression sizes created via CRISPR/Cas9-mediated targeted recombination. The candidate interval of *Cni-shls-1* locus is inferred by comparing two IR strains differing in introgression sizes and F1 phenotypes, i.e., the presence or absence of GFP+ F1 hybrids.

We reasoned that certain *C. nigoni*-specific alleles are key to the viability of hybrid progeny. To test this idea, we previously generated a large collection of *C. nigoni* strains, each carrying a GFP-linked *C. briggsae* genomic fragment as a heterozygous introgression (14, 17) (Fig. 1B, referred to as IR strain hereafter). Here we identified two *C. nigoni* genetic loci underlying the hybrid lethality by phenotyping F1 hybrids between individual IR strains and *C. briggsae* (Fig. 1B), because half of the cross progeny carry a homozygous *C. briggsae* introgression fragment, resulting in the absence of the syntenic *C. nigoni* genomic region. We first molecularly cloned a *C. nigoni*-derived hybrid lethality suppressor allele essential for hybrid embryo survival, and subsequently performed a genome-wide screen to map and clone another *C. nigoni*-specific allele responsible for the hybrid lethality. The former encodes a highly conserved <u>p</u>hospho<u>g</u>luco<u>m</u>utase (PGM), essential for glycosylation, while the latter encodes a *C. nigoni*-specific F-box protein involved in protein degradation. We demonstrate that the F-box gene specifically depletes both maternally and zygotically expressed *C. briggsae* PGM, but not *C. nigoni* PGM in the hybrid embryos. Finally, we show that the F-box gene was derived from a recent *C. nigoni*-specific duplication with rapid turnover even among populations, which causes unintended degradation of *C. briggsae* PGM in hybrids. Excessive expansion and fast evolution of F-box genes suggest their functions in immune response. Our findings delineate the inaugural mechanism underlying hybrid lethality between nematode species and reveal how recent gene duplication can establish reproductive barriers through unintended negative epistatic interactions.

## Results

### A hybrid lethality suppressor locus on *C. nigoni* chromosome IV (Chr. IV) is indispensable for hybrid F1 viability

We hypothesized that some specific *C. nigoni* alleles are essential for the viability of F1 and nB2 hybrids (Fig. 1A). To identify these hybrid lethality suppressor alleles, we previously conducted a systematic mapping of the essential *C. nigoni* genomic regions required for hybrid embryo viability using IR strains (see Methods for the generation of IR strains) (17). To achieve that, we performed reciprocal crosses between each IR strain and *C. briggsae* to produce F1 hybrids lacking specific *C. nigoni* genomic fragments (GFP-expressing ones) and examined lethal embryo phenotype in these hybrids (Fig. 1B). To facilitate phenotypic screening, we used the ratio of GFP-expressing hybrid F1 adults as a proxy for the proportion of lethal embryos. This is because hybrid embryo lethality was complicated by the fact that over half of the F1 hybrids from wild type (WT) cross arrested as embryo (13, 15) (see Methods for phenotyping of hybrid progeny from IR strains and *C. briggsae*).

The mapping allows us to identify ZZY10018, an IR strain carrying a ∼4.7Mb fragment from the right arm of *C. briggsae* Chr. IV, that barely produced any GFP-expressing F1 hybrid adults with *C. briggsae* regardless of parent of origin (IR as mother: 0%; IR as father: 0.6%). This was significantly deviating from the expected ratio of 50% of GFP-expressing F1 hybrid adults (*P*<0.0001, Chi-square test) (Fig. 1C and D). Consistently, the hatching rates of F1 hybrids decreased approximately by half with ZZY10018 as the mother (30.8%) or father (22.6%) compared to WT crosses (*C. nigoni* as mother: 55.4%; *C. nigoni* as father: 41.0%, Fig. S1A). Such decrease was mainly due to the lethality of GFP-expressing embryos because the ratio of arrested F1 hybrid embryos expressing GFP was approximately 70%, significantly higher than the expected ratio of 50% (*P*<0.0001, Chi-square test, Fig. S1B). Taken together, these results indicate that the absence of a *C. nigoni* locus on *C. nigoni* Chr. IV led to complete embryonic lethality in F1 hybrids. We therefore named this locus as <u>s</u>pecies <u>h</u>ybrid lethality <u>s</u>uppressor (*Cni-shls-1*).

To map the candidate interval of *Cni-shls-1* locus, we generated IR strains carrying various sizes of introgression fragments from the right arm of *C. briggsae* Chr. IV and characterized the phenotypes of their F1 hybrids with *C. briggsae* (Fig. 1E and Fig. 2A). To facilitate recombination during crossing, we generated IR strains with desirable introgression sizes using a recently developed targeted recombination method mediated by CRISPR/Cas9 (18) (See Method for targeted recombination, Fig. 1E).

**Figure 2.**
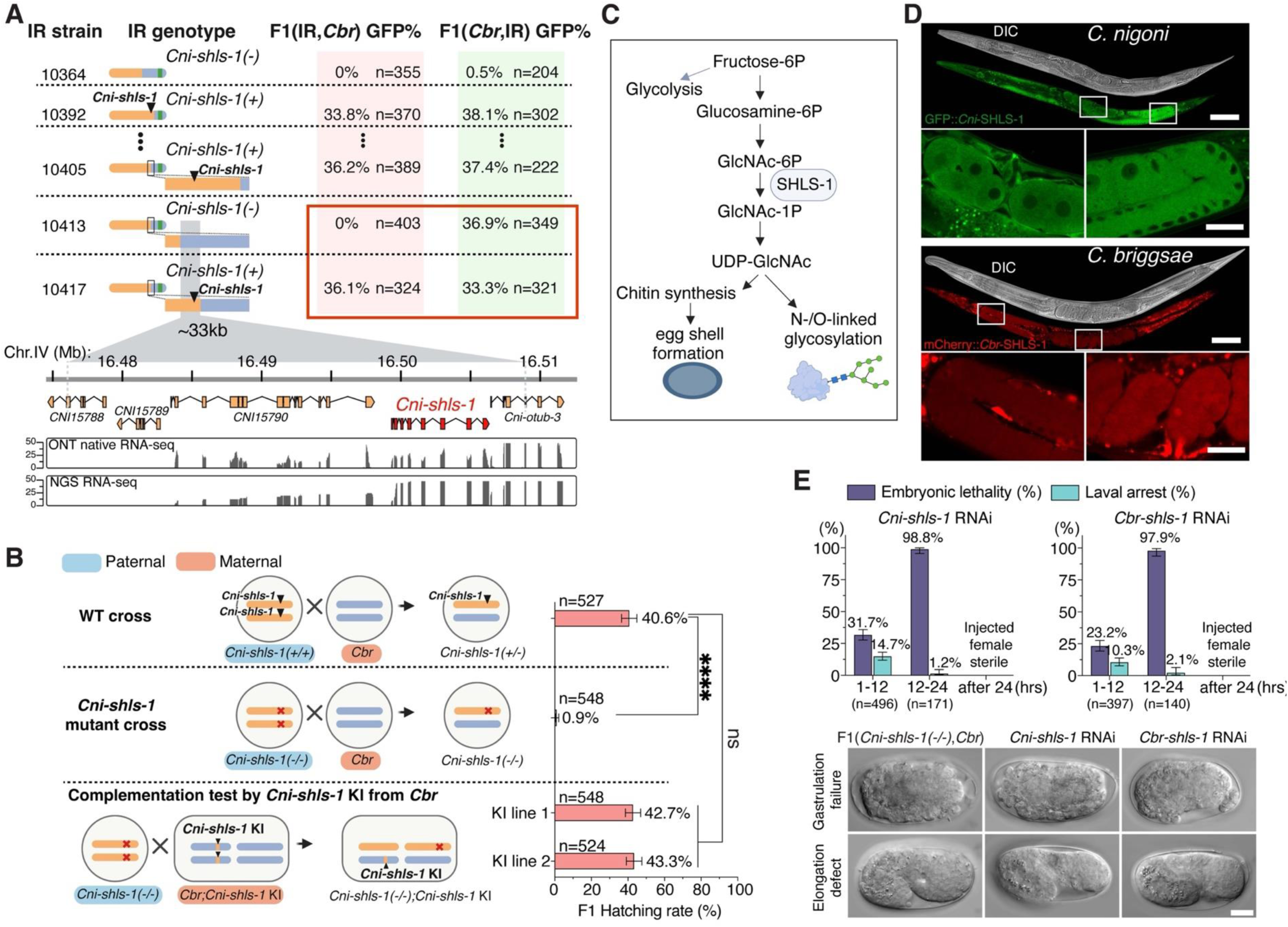
*Cni-shls-1* encodes a conserved and essential phosphoglucomutase. **(A)** Top: Fine-mapping of the *Cni-shls-1* locus using IR strains with varying *C. briggsae* introgression sizes. Percentages of GFP+ hybrid F1 adults from *C. briggsae* mothers or fathers are shaded with light pink and light green, respectively. The genotypes, i.e., *Cni-*Chr. IVs with various *C. briggsae* introgression sizes, of the IR strains are shown. Bottom: Predicted gene models within the ∼33kb candidate interval. RNA-seq read coverage (ONT and NGS) for *C. nigoni* embryos is shown below. A red rectangle highlights the interval responsible for phenotypic change (presence/absence of GFP+ F1 hybrids) from *C. briggsae* mothers, not fathers. **(B)** Validation of *Cni-shls-1* as a hybrid lethality suppressor gene. Shown is the comparison of hatching rates among F1 hybrids from WT parents (top), *Cni-shls-1(-/-)* mutant fathers with *C. briggsae* mothers (middle), and *Cni-shls-1(-/-)* mutant fathers with *C. briggsae* mothers carrying a *Cni-shls-1* knock-in (KI) (bottom). Genotype schematics for parents and F1 hybrids from each cross are shown on the left. \*\*\*\**P*<0.0001, Fisher’s exact test; error bars: 95% confidence interval. **(C)** Hexosamine biosynthesis pathway in nematodes. SHLS-1 is a phosphoglucomutase converting GlcNAc-6P to GlcNAc-1P. Fructose-6P: Fructose 6-phosphate; Glucosamine-6P: Glucosamine 6-phosphate; GlcNAc-6P: N-acetylglucosamine-6-phosphate; GlcNAc-1P: N-acetylglucosamine-1-phosphate; UDP-GlcNAc: Uridine diphosphate N-acetylglucosamine. **(D)** Expression of SHLS-1 in adult female *C. nigoni* (top) and *C. briggsae* (bottom), shown with N-terminally tagged GFP::*Cni-*SHLS-1 and mCherry::*Cbr-*SHLS-1, respectively. Magnified images highlight expression in early embryos and germlines. Scale bar: 100µm (original) and 20µm (magnified). **(E)** Reduced *shls-1* expression causes various defects, including embryonic lethality in both *C. nigoni* and *C. briggsae*. Top: Percentages of embryonic lethality and larval arrest in progeny from two 12-hour intervals post-RNAi against *shls-1* in *C. nigoni* (left) and *C. briggsae* (right). Bottom: Representative images of arrested embryos with gastrulation failure (top) or elongation defects (bottom) in F1 hybrids from *Cni-shls-1(-/-)* fathers and *C. briggsae* mothers (left), and in *shls-1* RNAi-treated *C. nigoni* (middle) and *C. briggsae* (right). Scale bar, 10µm.

### *Cni-shls-1* encodes a conserved and essential phosphoglucomutase

The *Cni-shls-1* locus was initially mapped to an interval of roughly 3Mb by comparing two IR strains with different introgression fragment sizes (ZZY10364: ∼4.01Mb and ZZY10392: ∼1.23Mb) (Fig. 2A). ZZY10364 produced nearly no GFP-expressing hybrid F1 adults with *C. briggsae* mothers (0%) or fathers (0.5%), whereas over 30% GFP-expressing hybrid F1 adults were observed following reciprocal crosses between ZZY10392 and *C. briggsae* (Fig. 2A). The interval was further narrowed using a series of IR strains with various *C. briggsae* Chr. IV introgression sizes (Fig. 2A; see Table. S1 for a full list of the IR strains used). Based on the absence and presence of GFP-expressing adult F1 hybrids derived from the two IR strains ZZY10413 (0%) and ZZY10417 (36.1%) as the fathers, we finally narrowed the candidate interval to ∼33 kb (Fig. 2A). Surprisingly, reciprocal crosses of both strains with *C. briggsae* fathers demonstrated no differential phenotypes (36.9% vs. 33.3%, *P* = 0.4095; Fisher’s exact test, Fig. 2A), indicating that at least two independent hybrid lethality suppressor loci exist on *C. nigoni* Chr. IV. For all subsequent experiments, we focused on the *Cni-shls-1* locus that produces F1 progeny survival in the cross between IR strain fathers and *C. briggsae* mothers.

To establish the molecular identity of *Cni-shls-1*, we first examined the four candidate genes within the ∼33kb interval (Fig. 2A). To confirm the predicted gene models, we performed Oxford Nanopore native-RNA sequencing (RNA-seq) and aligned the sequencing reads along with existing Illumina short RNA-seq reads from *C. nigoni* embryos to the interval. Only the gene models of *CNI15790* and *CNI15791* were highly supported (Fig. 2A). We next performed simultaneous and individual deletion of *CNI15790* and *CNI15791* on the *C. nigoni* Chr. IV associated with the *C. briggsae* introgression in ZZY10417. No GFP-expressing hybrid F1 adults were observed when both genes or only *CNI15791* were deleted (Fig. S2), indicating that *CNI15791* is *Cni-shls-1*.

To evaluate whether *Cni-shls-1* functions independently of the associated *C. briggsae* Chr. IV introgression, we next deleted *Cni-shls-1* in WT *C. nigoni* and crossed the homozygous mutant males with *C. briggsae* females. Nearly all F1 hybrids failed to hatch (99.1%), and the rare escapers arrested at the L1 larval stage. This is in sharp contrast to the ratio of hatched hybrid F1 progeny from the WT cross (40.6%, *P*<0.0001, Fisher’s exact test, Fig. 2B). To further confirm that the embryonic lethality was caused by the loss of *Cni-shls-1*, we performed a complementation test. We first expressed *Cni-shls-1* as an extrachromosomal array either in *Cni-shls-1(-/-)* mutant fathers or *C. briggsae* mothers. The complete embryonic lethality could be partially rescued when *Cni-shls-1* was expressed in the F1 hybrids through the array inherited from either parent, using two independent array lines (array from *Cni-shls-1(-/-)* fathers: 18.9% and 13.8%; array from *C. briggsae* mothers: 16.2% and 16.1%, vs. 0.9%, *P*<0.0001, Fisher’s exact test, Fig. S3). The partial rescue appeared to result from the mosaicism of transgenic animals, because all the hatched animals carried the *Cni-shls-1* array indicated by the presence of injection marker (Fig. S3). Full rescue of F1 hybrids was achieved by a single copy knock-in of *Cni-shls-1* inherited from *C. briggsae* mothers using two independent *C. briggsae* knock-in lines (42.7% and 43.3% vs 0.9%, *P*<0.0001, Fisher’s exact test, Fig. 2B). These results support that the embryonic lethality was suppressed when *Cni-shls-1* was present.

To our surprise, instead of a fast-evolving gene, *Cni-shls-1* encodes a highly conserved phosphoglucomutase (PGM), orthologous to PGM3 in humans (Fig. 2C). PGM catalyzes the conversion of N-acetylglucosamine-6-phosphate to N-acetylglucosamine-1-phosphate in the hexosamine pathway (Fig. 2C). This essential metabolic pathway, conserved across species, converts glucose to uridine diphosphate N-acetylglucosamine (UDP-GlcNAc), the key substrate for O- and N-linked glycosylation and chitin synthesis (19). We tested whether the *shls-1* orthologues were essential in *C. nigoni* and *C. briggsae*. Indeed, although *Cni-shls-1(-/-)* males were fertile, the homozygous mutant females were completely sterile (Fig. S4). We generated a *Cbr-shls-1* deletion mutant and observed a similar phenotype (Fig. S4). Oocyte development appeared normal, but fertilized eggs in both *Cni-shls-1(-/-)* and *Cbr-shls-1(-/-)* mutants were severely malformed, reminiscent of eggshell formation defects from perturbation of *C. elegans* chitin synthase genes (20, 21) (Fig. S4). These results suggest a disruption of hexosamine pathway in *shls-1(-/-)* mutant females, leading to UDP-GlcNAc deficiency and subsequent sterility.

However, the viability of *shls-1(-/-)* mutants implies maternal loading of SHLS-1 from the heterozygous mutants allows embryogenesis. To test this, we tagged GFP and mCherry to the N-terminus of endogenous *Cni-*SHLS-1 and *Cbr-*SHLS-1, respectively, and examined their expression patterns. Both *gfp::Cni-shls-1* and *mCherry::Cbr-shls-1* animals were homozygous viable with no obvious defects, indicating fully functional knock-in alleles. These proteins were ubiquitously expressed in almost all cells, enriched in gonads of both sexes, and in early embryos of both species (Fig. 2D and Fig. S5), supporting a maternal loading of the enzyme. We further examined whether a decreased SHLS-1 dosage led to the embryogenesis defects. To this end, we knocked down *shls-1* by injecting double-stranded RNAs into the gonads of both species. In the first 12 hours post injection, each species produced arrested embryos (*Cni*: 31.7%; *Cbr:* 23.2%) and larvae (*Cni:* 14.7%; *Cbr:* 10.3%) (Fig. 2E). The phenotypes then turned more severe such that much fewer embryos were laid 12-24 hours post injection, and nearly all failed to hatch (*Cni:* 98.8%; *Cbr:* 97.9%, Fig. 2E). No more embryos were laid beyond 24 hours, resembling the *shls-1(-/-)* mutant females (Fig. 2E). Interestingly, we observed similar terminal phenotypes, including failures in gastrulation and elongation of the arrested embryos for *shls-1* perturbed progeny laid in the first 12 hours and hybrid F1 progeny from *Cni-shls-1(-/-)* mutant fathers (Fig. 2E). Collectively, these results demonstrate a highly conserved and essential role of SHLS-1 in both *C. briggsae* and *C. nigoni*. Furthermore, the embryonic lethality in the F1 hybrids between *C. briggsae* mothers and *Cni-shls-1(-/-)* fathers appears to result from the deficiency of SHLS-1 enzymes.

### Specific and rapid degradation of *C. briggsae* SHLS-1 in F1 hybrids during early embryogenesis

Having established that *shls-1* is functionally conserved in both species, we wondered why *Cni-shls-1* is indispensable for the survival of their F1 hybrids. We examined sequence variations of SHLS-1 orthologues across nine nematodes. As expected, the proteins were highly conserved, even between more distantly related species such as *Pristionchus pacificus* and *Oscheius tipulae* (Fig. S6), indicative of purifying selection acting on these proteins. In agreement with this, although there were 36 single amino acid changes from the pairwise protein sequence alignment between *Cni-*SHLS-1 and *Cbr-*SHLS-1, none included the predicted active sites of the enzymes (Fig. 3A and Fig. S7A), and the only divergent motif between the two is not responsible for their differential roles for F1 hybrid viability (Fig. 3A and Fig. S7B and C). Comparison of tertiary structures also reveals high conservation among the two SHLS-1 and even human PGM3 (Fig. 3B). Taken together, these results demonstrate extensive functional conservation between *Cni-*SHLS-1 and *Cbr-*SHLS-1.

**Figure 3.**
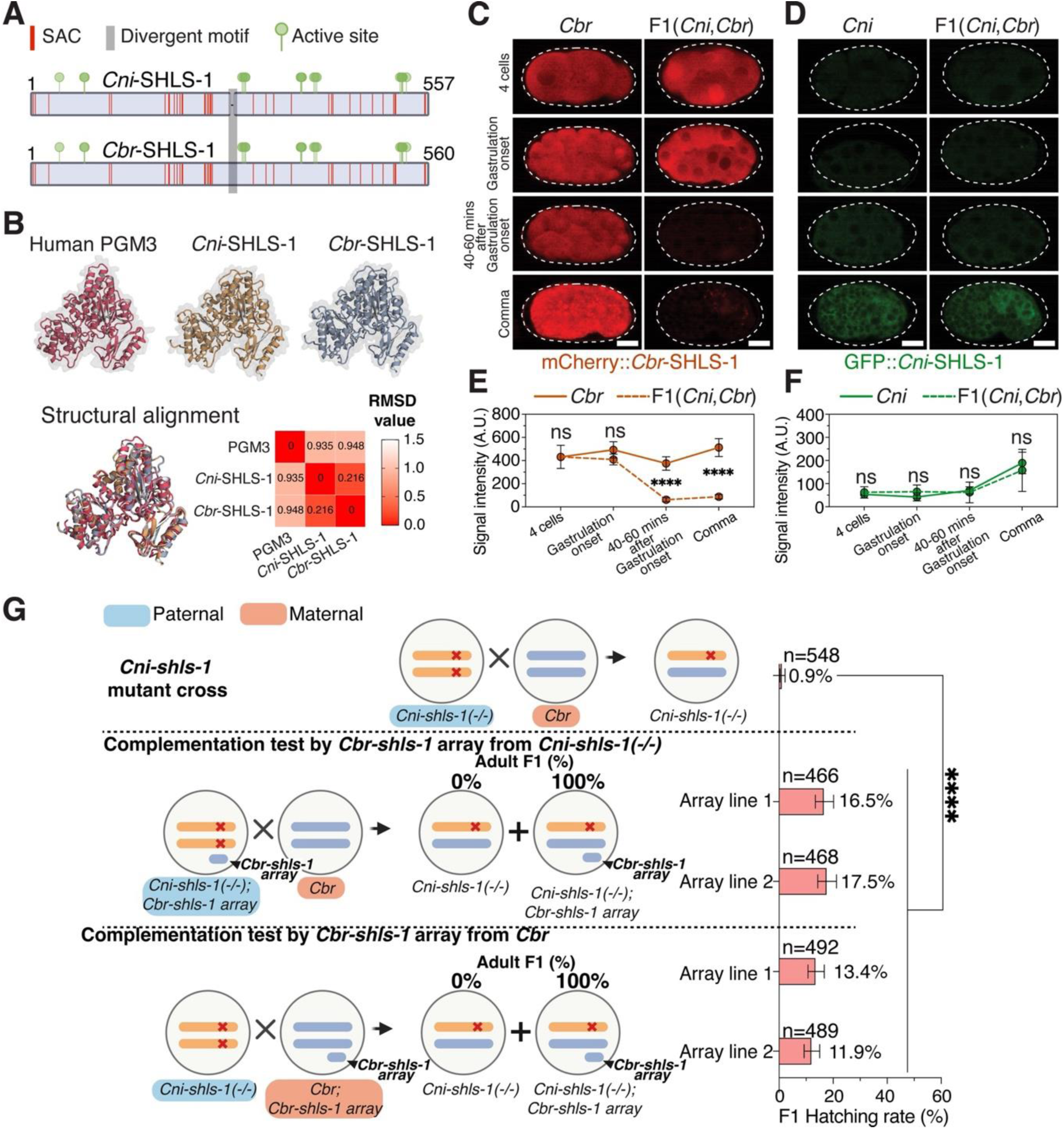
Specific degradation of *C. briggsae* SHLS-1 in F1 hybrids during early embryogenesis. **(A)** Protein sequence alignment of *Cni-*SHLS-1 and *Cbr-*SHLS-1. The single amino acid change (SAC), divergent motif and active sites of the two proteins are indicated. **(B)** Top: comparison of Alphafold2-predicted protein structures for human PGM3, *Cni-*SHLS-1 and *Cbr-*SHLS-1. Bottom: Structural alignment of the three proteins with corresponding RMSD values. **(C-D)** Rapid degradation of *C. briggsae* SHLS-1 in F1 hybrids. Representative images show *Cbr-*SHLS-1 (C) and *Cni-*SHLS-1 (D) expression in embryos of *C. briggsae* carrying heterozygous *mCherry::Cbr-shls-*1, *C. nigoni* carrying heterozygous *gfp::Cni-shls-1*, and F1 hybrids from *C. nigoni* fathers carrying homozygous *gfp::Cni-shls-1* and *C. briggsae* mothers carrying homozygous *mCherry::Cbr-shls-1*, across four developmental stages: 4-cell, gastrulation onset, 40-60 mins after gastrulation onset and comma stage. Genotype details: *Cbr* embryos are *mCherry::Cbr-shls-1(+/-)* from WT fathers and *mCherry::Cbr-shls-1(+/+)* mothers; *Cni* embryos are *gfp::Cni-shls-1(+/-)* from WT mothers and *gfp::Cni-shls-1(+/+)* fathers; *F1(Cni,Cbr)* embryos are *mCherry::Cbr-shls-1(+)/gfp::Cni-shls-1(+)* from *mCherry::Cbr-shls-1(+/+)* mothers and *gfp::Cni-shls-1(+/+)* fathers. Scale bar, 10µm. **(E-F)** Quantification of mCherry (E) or GFP intensity (F) in hybrid F1 embryos from *C. nigoni* fathers and *C. briggsae* mothers compared to that in *C. briggsae* (mCherry intensity) or *C. nigoni* embryos (GFP intensity) at each stage shown in C and D. n=5 for each stage; \*\*\*\**P<*0.0001, unpaired t-test; Error bar: s.e.m; A.U.: Arbitrary units. **(G)** Rescuing F1 hybrid lethality via *Cbr-shls-1* overexpression. Shown is the comparison of hatching rates for F1 hybrids from *Cni-shls-1(-/-)* fathers and *C. briggsae* mothers (top), and hybrids carrying a *Cbr-shls-1* extrachromosomal array inherited from either the *Cni-shls-1(-/-)* fathers (middle) or the *C. briggsae* mothers (bottom). Genotype schematics (Chr. IV only) for parents and F1 hybrids are shown on the left. Also shown are the percentages of adult F1 hybrids with/without the array. Two independent array-bearing lines were used for each complementation cross. \*\*\*\**P*<0.0001, Fisher’s exact test; error bars: 95% confidence interval.

We next asked whether the expression of *Cbr-*SHLS-1 was selectively perturbed in F1 hybrid embryos. We observed comparable expression patterns of either transgene (*mCherry::Cbr-shls-1* or *gfp::Cni-shls-1* as shown in Fig. 2D) as heterozygote in *C. briggsae* and *C. nigoni* embryos: When the transgenic animals carrying *mCherry::Cbr-shls-1(+/+)* or *gfp::Cni-shls-1(+/+)* were used as the mothers and crossed with a WT fathers, SHLS-1 showed bright expression throughout embryogenesis due to both maternal supply and zygotic expression (Fig. 3C; Fig. S8A; Fig. S9B; Movie 2 and 6). In the reciprocal cross, we only observed SHLS-1 expression in late embryos (Fig. 3D; Fig. S8B; Fig. S9A; Movie 3 and 5), indicating zygotic expression.

We then examined the expression dynamics of *Cni-*SHLS-1 and *Cbr-*SHLS-1 in the F1 hybrids from the cross between *C. briggsae mCherry::Cbr-shls-1(+/+)* mothers and *C. nigoni gfp::Cni-shls-1(+/+)* fathers. Strikingly, we observed a rapid depletion of the maternally deposited *Cbr-*SHLS-1 in hybrid embryos starting from approximately 26-cell stage (Fig. 3C; Fig. S8C; Movie 1). Around 40-60 minutes after the onset of gastrulation, the expression intensity of *Cbr-*SHLS-1 dropped to background levels (Fig. 3E; Movie 1 and 2). In contrast, the intensity of zygotically expressed *Cni-*SHLS-1 was comparable to that of the heterozygous WT control (Fig. 3D and F; Fig. S9; Movie 1 and 3). Similar degradation of zygotically expressed *Cbr-*SHLS-1 was observed in F1 hybrids from the reciprocal cross (Fig. S9A, C and D; Movie 4 and 5). As expected, no degradation of maternally deposited *Cni-*SHLS-1 was observed in F1 hybrid embryos when compared with the heterozygous control (Fig. S9B, C and E; Movie 4, and 6). These results clearly demonstrate that the embryonic lethality of the F1 hybrids lacking *Cni-shls-1* stems from specific degradation of *Cbr-*SHLS-1. To further confirm this, we overexpressed *Cbr-shls-1* from an extrachromosomal array in the F1 hybrids and quantified their hatching rate. The lethality was partially rescued when the *Cbr-shls-1* array was inherited from either the *Cni-shls-1(-/-)* mutant fathers (16.5% and 17.5% vs. 0.9%, *P*<0.0001, Fisher’s exact test) or the *C. briggsae* mothers (13.4% and 11.9% vs. 0.9%, *P*<0.0001, Fisher’s exact test) (Fig. 3G), supporting that *Cbr-*SHLS-1 degradation led to hybrid embryonic lethality.

### A *C. nigoni*-specific gene causes hybrid lethality through selective depletion of *Cbr-*SHLS-1

We next asked why *Cbr-*SHLS-1 was selectively degraded in the F1 hybrids between the two species. Although hybrid incompatibility in intraspecific nematodes often follows a toxin-antidote (TA) model (22–25), they usually involve species-specific gene pairs, which is not the case with *shls-1*. Instead, our results align well with the Bateson-<u>D</u>obzhansky-<u>M</u>uller incompatibility (DMI) model (4), which posits that HI typically arises from improper interactions between at least two genes from different parental species. While each gene is innocuous in its native genomic context, the incompatibility manifests when they encounter in hybrids. Therefore, we hypothesized that another *C. nigoni* locus was responsible for the specific *Cbr-*SHLS-1 depletion (Fig. 4A). We named this locus *Cni-neib-1* (*C. nigoni* <u>n</u>egative <u>e</u>pistatic interactor with *C<u>b</u>r-shls-1*). To test this hypothesis, we performed a genome-wide screening to map and clone the *Cni-neib-1* locus. We reasoned that, in contrast to the F1 hybrid death in the absence of *Cni-shls-1*, the additional loss of another *C. nigoni* genomic region containing *Cni-neib-1* is expected to rescue the lethality (Fig. 4B).

**Figure 4.**
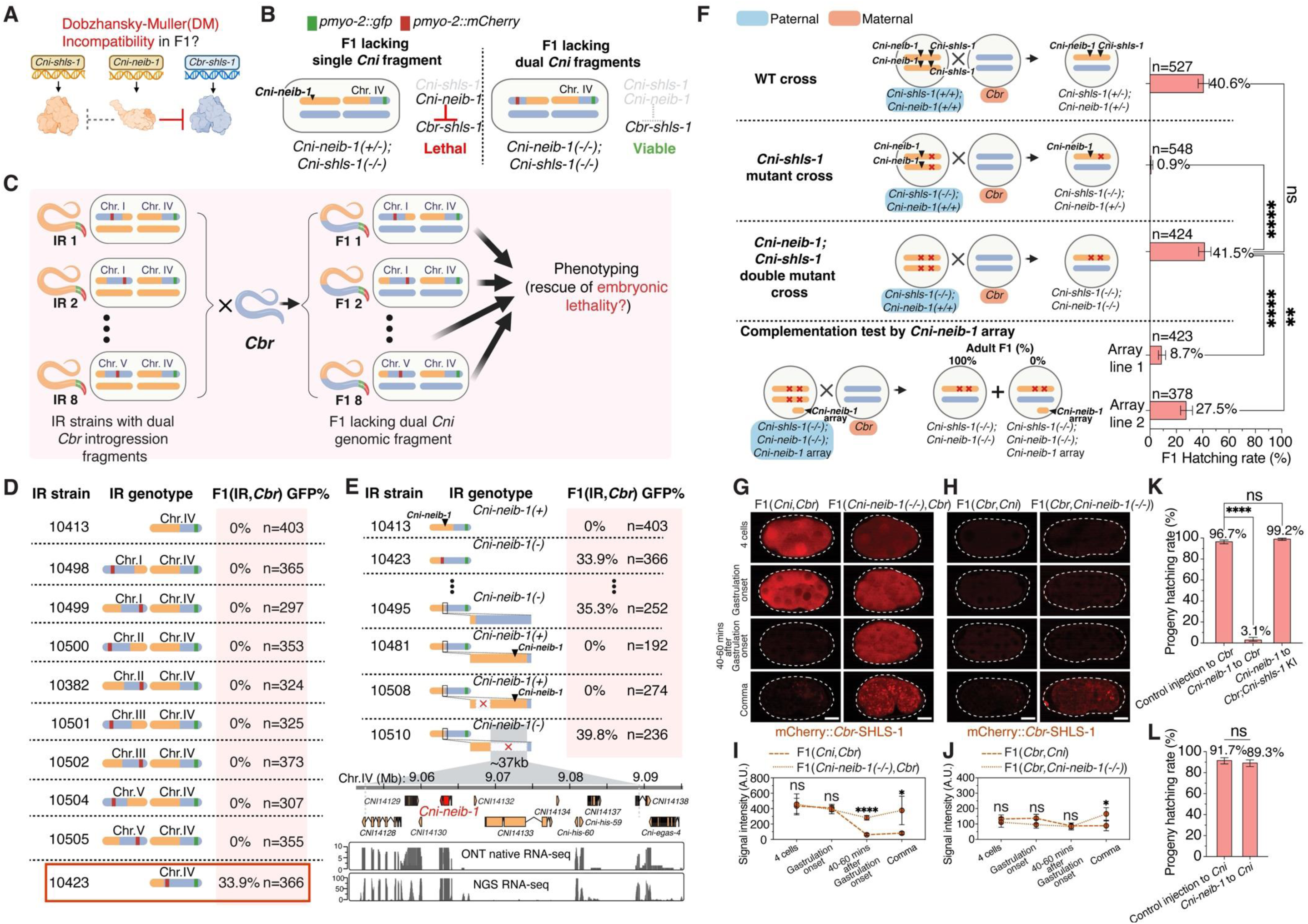
The hybrid lethality results from a two-gene incompatibility between *C. briggsae shls-1* and *C. nigoni neib-1*. **(A)** Schematic illustrating negative epistatic interaction between *Cni-neib-1* and *Cbr-shls-1*, but not *Cni-shls-1*. This interaction explains the F1 hybrid lethality, consistent with the Dobzhansky-Muller (DM) incompatibility model. **(B)** Schematic illustrating the rationale for rescuing hybrid lethality in F1 hybrids lacking dual *C. nigoni* genomic fragments. *Cni-neib-1* negatively interact with *Cbr-shls-1* in the F1 hybrids lacking *C. nigoni* Chr. IV right arm containing *Cni-shls-1* (left). Additional absence of another *C. nigoni* fragment containing *Cni-neib-1* is expected to rescue the lethality (right). **(C)** Schematic of the screening strategy for *Cni-neib-1* by phenotyping F1 hybrids lacking dual *C. nigoni* genomic fragments. IR strains carrying dual *C. briggsae* introgression fragments were first generated through by crossing ZZY10413 (*C. briggsae* Chr. IV right arm introgression, see Fig. 2A) with another IR strain carrying the left or the right half of a *C. briggsae* autosome (mCherry-labeled). Each IR strain was subsequently crossed with *C. briggsae* mothers, and hybrid progeny homozygous for both introgressions were phenotyped for GFP+ F1 hybrids. **(D)** Rescuing hybrid lethality in F1 hybrids with homozygosity for an extended *C. briggsae* Chr. IV introgression. Shown is the comparison of percentages of GFP+ hybrid F1 adults fathered by IR strains carrying *C. briggsae* Chr. IV right arm (ZZY10413), both *C. briggsae* Chr. IV right arm and either half of another autosome, or an extended *C. briggsae* Chr. IV (highlighted by a red rectangle). Genotype schematics for *C. nigoni* chromosomes with various *C. briggsae* introgression sizes in the IR strains are shown. **(E)** Top: Fine-mapping of *Cni-neib-1* locus by comparing IR strains with varying *C. briggsae* introgression sizes or large genomic deletions on *C. nigoni* chromosomes with introgressions. Percentages of GFP+ hybrid F1 adults fathered by the IR strains with corresponding genotypes i.e., *Cni-* Chr. IVs with various *C. briggsae* introgressions, are shown. Bottom: Predicted gene models within the ∼37kb candidate interval. RNA-seq read coverage (ONT and NGS) for *C. nigoni* embryos is shown below. **(F)** Validation of *Cni-neib-1*. Shown is the comparison of hatching rates for F1 hybrids from WT parents (top, see Fig. 2B), F1 hybrids from *Cni-shls-1(-/-)* mutant fathers and *C. briggsae* mothers (middle-top, see Fig. 2B), F1 hybrids from *Cni-shls-1(-/-);Cni-neib-1(-/-)* double mutant fathers and *C. briggsae* mothers (middle-bottom), and F1 hybrids carrying a *Cni-neib-1* extrachromosomal array inherited from the *C. nigoni* double mutant fathers (bottom). Genotype Schematics (Chr. IV only) for parents and F1 hybrids are shown on the left. Also shown are the percentages of adult F1 hybrids with/without the array. Two independent *Cni-neib-1* array-bearing lines were used for complementation. \*\*\*\**P*<0.0001, \*\**P*=0.0041, Fisher’s exact test; error bars: 95% confidence interval. **(G-H)** Deletion of *Cni-neib-1* prevents degradation of *Cbr-*SHLS-1 in F1 hybrids. Representative images of *Cbr-*SHLS-1 expression in hybrid F1 embryos from *C. briggsae* carrying homozygous *mCherry::Cbr-shls-1* as the mothers (G) or fathers (H) across the same developmental stages shown in Fig 3C and D. The *mCherry::Cbr-shls-1(+/+) C. briggsae* was crossed with either *C. nigoni* (left) or *Cni-neib-1(-/-)* mutant (right) in both directions. Scale bar, 10µm. **(I-J)** Quantification of mCherry signal intensity in F1 hybrid embryos from *C. nigoni* or *Cni-neib-1(-/-)* mutant at each stage shown in G and H. F1 hybrid embryos were obtained from reciprocal crosses with *mCherry::Cbr-shls-1(+/+) C. briggsae* as mothers (I) or fathers (J). n=5 for each stage; \*\*\*\**P*<0.0001, * *P*<0.05, unpaired t-test. Error bar: s.e.m. A.U., Arbitrary units. **(K-L)** Introduction of *Cni-neib-1* genomic fragments remarkably decreases hatching rates in *C. briggsae* but not *C. nigoni*. Shown is the comparison of hatching rates of the progeny from *C. briggsae* (K) or *C. nigoni* (L) injected with control buffer or *Cni-neib-1* genomic DNA. *Cbr;Cni-shls-1* KI worm is the transgenic *C. briggsae* carrying a *Cni-shls-1* knock-in (see Fig. 2B). Note that ectopic *Cni-shls-1* expression in *C. briggsae* rescues the embryonic lethality. \*\*\*\**P*<0.0001, Fisher’s exact test. Error bars: 95% confidence intervals.

To generate F1 hybrids lacking both *Cni-neib-1* and *Cni-shls-1* loci, we first generated eight IR strains carrying dual *C. briggsae* introgressions, with one derived from either half of a *C. briggsae* autosome excluding Chr. IV (mCherry-labeled), and the other derived from Chr. IV right arm (GFP-labeled) (see Methods, Fig. 4C and Fig. S10). We then crossed each dual-introgression IR strain with *C. briggsae* mothers to assess whether hybrid lethality could be rescued, indicated by the presence of fluorescent marker-labeled hybrid F1 adults (Fig. 4C). Chr. X was excluded because most IR strains carrying *C. briggsae* Chr. X introgression are sterile or inviable, incapable of crossing (14). However, we observed no GFP-expressing F1 hybrids across all crosses from the eight dual-introgression IR strain fathers (Fig. 4D), suggesting that *Cni-neib-1* is linked with *Cni-shls-1* on Chr. IV (Fig. S11). Indeed, an IR strain (ZZY10423) with a substantially longer *C. briggsae* Chr. IV introgression fragment rescued the hybrid F1 lethality (Fig. 4D).

To map the *Cni-neib-1* locus, we again generated IR strains with various lengths of *C. briggsae* Chr. IV introgressions (Fig. 4E). The target interval was initially mapped to ∼53 kb, and subsequently narrowed to ∼37kb by large genomic deletions (Fig. 4E). We then aligned both Oxford nanopore native RNA-seq and Illumina short RNA-seq reads from *C. nigoni* embryos to the region and validated the gene models within the interval. Among the ten predicted genes within the interval, we prioritized for subsequent characterization the three genes with the most abundant expression (Fig. 4E), i.e., *CNI14129*, *CNI14130, CNI14131*. Simultaneous deletion of *CNI14129* and *CNI14130* on the *C. nigoni* Chr. IV associated with the *C. briggsae* introgression in ZZY10481 failed to rescue the hybrid F1 lethality, whereas the deletion of *CNI14131* rescued the hybrid embryo viability (Fig. S12), confirming that *CNI14131* is *Cni-neib-1*.

To avoid possible complication associated with the introgression segment, we generated a *Cni-neib-1(-/-)*, *Cni-shls-1(-/-)* double knockout mutant in WT *C. nigoni* males. Hatching rates of their F1 hybrids with *C. briggsae* females was significantly higher than that from *Cni-shls-1(-/-)* single mutant fathers (41.5% vs. 0.9%, *P*<0.0001, Fisher’s exact test, Fig. 4F), at a level comparable to that of the WT cross (40.6%, Fig. 4F). The rescue could also be reversed when a *Cni-neib-1* extrachromosomal array inherited from the double mutant fathers was crossed to the F1 hybrids. Specifically, hatching rates dropped to 8.7% and 27.5% for F1 hybrids fathered by two independent *Cni-neib-1* array bearing lines, respectively, and all hatched progeny carried no *Cni-neib-1* array as indicated by the injection marker (Fig. 4F), supporting a strong negative interaction between *Cni-neib-1* and *Cbr-shls-1* in hybrids.

To confirm that *Cni-neib-1* leads to specific degradation of *Cbr-*SHLS-1, we generated a single *Cni-neib-1* deletion allele in the WT *C. nigoni*. The homozygous mutant was viable with no obvious defects. We then crossed the mutant with *C. briggsae* transgenic animals carrying *mCherry::Cbr-shls-1(+/+)* in reciprocal directions and examined *Cbr-*SHLS-1 expression in hybrid F1 embryos. Consistent with the hybrid lethality rescue, *Cbr-*SHLS-1 expression intensity remained constantly high in hybrid F1 embryos fathered by a *Cni-neib-1(-/-)* mutant compared to a cross using *Cni-neib-1(+/+)* animals (Fig. 4G and I, Fig. S13A and B; Movie 1 and 7). *Cbr-*SHLS-1 expression was also detected at the late embryonic stages in F1 hybrids when *Cni-neib-1(-/-)* was the mother (Fig. 4H and J; Fig. S13C and D, Movie 4 and 8). Collectively, these results indicate that the absence of *Cni-neib-1* abolished the degradation of both maternally and zygotically expressed *Cbr-*SHLS-1.

Finally, we directly examined the capacity of *Cni-*NEIB-1 to deplete *Cbr-*SHLS-1 in WT *C. briggsae.* To this end, we amplified *Cni-neib-1* genomic fragments spanning the putative promoter to the 3’ UTR region and injected the DNA into *C. briggsae*. Strikingly, we observed a significantly lower hatching rate in the resulting progeny (3.1%), in contrast to the background level in *C. briggsae* injected with control buffer (96.7%, *P*<0.0001, Fisher’s exact test, Fig. 4K). Embryonic development was fully rescued (99.2% hatching) when injecting the same *Cni-neib-1* DNA into transgenic *C. briggsae* carrying an *Cni-shls-1* knock-in copy as shown in Fig. 2B (Fig. 4K). The lethality was unlikely to be caused by overdosage, as injecting the same *Cni-neib-1* DNA into *C. nigoni* did not affect progeny viability (Fig. 4L). Taken together, we demonstrate that *Cni-neib-1* specifically depleted *Cbr-*SHLS-1 protein in both hybrids and native *C. briggsae* species, leading to embryonic lethality.

### *Cni-neib-1* is a recently duplicated F-box gene in *C. nigoni*

To understand the molecular mechanism and evolutionary trajectory of the specific depletion of *Cbr-*SHLS-1, we examined the conservation of *Cni-neib-1* across nematode species. In contrast to *shls-1*, *Cni-neib-1* encodes a *C. nigoni*-specific F-box gene. F-box proteins function as essential adaptors for specific substrate binding in the SCF (Skp, Cullin, F-box) complex, one of the E3 ubiquitin ligases involved in protein ubiquitination and subsequent degradation (26) (Fig. 5A). These proteins typically feature a conserved F-box domain and a divergent substrate-binding domain which targets specific protein for degradation. *Cni-*NEIB-1 contains an F-box domain and an F-box-associated type 2 (FBA2) domain at the N- and C terminus, respectively (Fig. 5A). F-box genes are commonly expanded in *Caenorhabditis* nematodes (11, 27). Interestingly, our examination of protein families significantly enriched in *C. nigoni* compared to *C. briggsae* revealed that F-box proteins are the most expanded. (Fig. 5B, Fig. S14). *Cni-neib-1* arose from a very recent gene duplication, which produced 33 paralogous copies that are unique to the *C. nigoni* genome (Table S2). The most closely related F-box proteins in *C. briggsae* (*CBG13797, CBG19758* and *CBG27620*) and *C. elegans* (*Y45F10B.12* and *T02G6.11*) share only ∼20% identity in protein sequence with *Cni-*NEIB-1 (Fig. 5C), whereas identity among *the C. nigoni* paralogues is at least 40% (Fig. 5C), supporting a recent F-box gene expansion exclusively in *C. nigoni*. We therefore named the protein family “F-box of *Cni-neib-1* (*fbxn*)” family.

**Figure 5.**
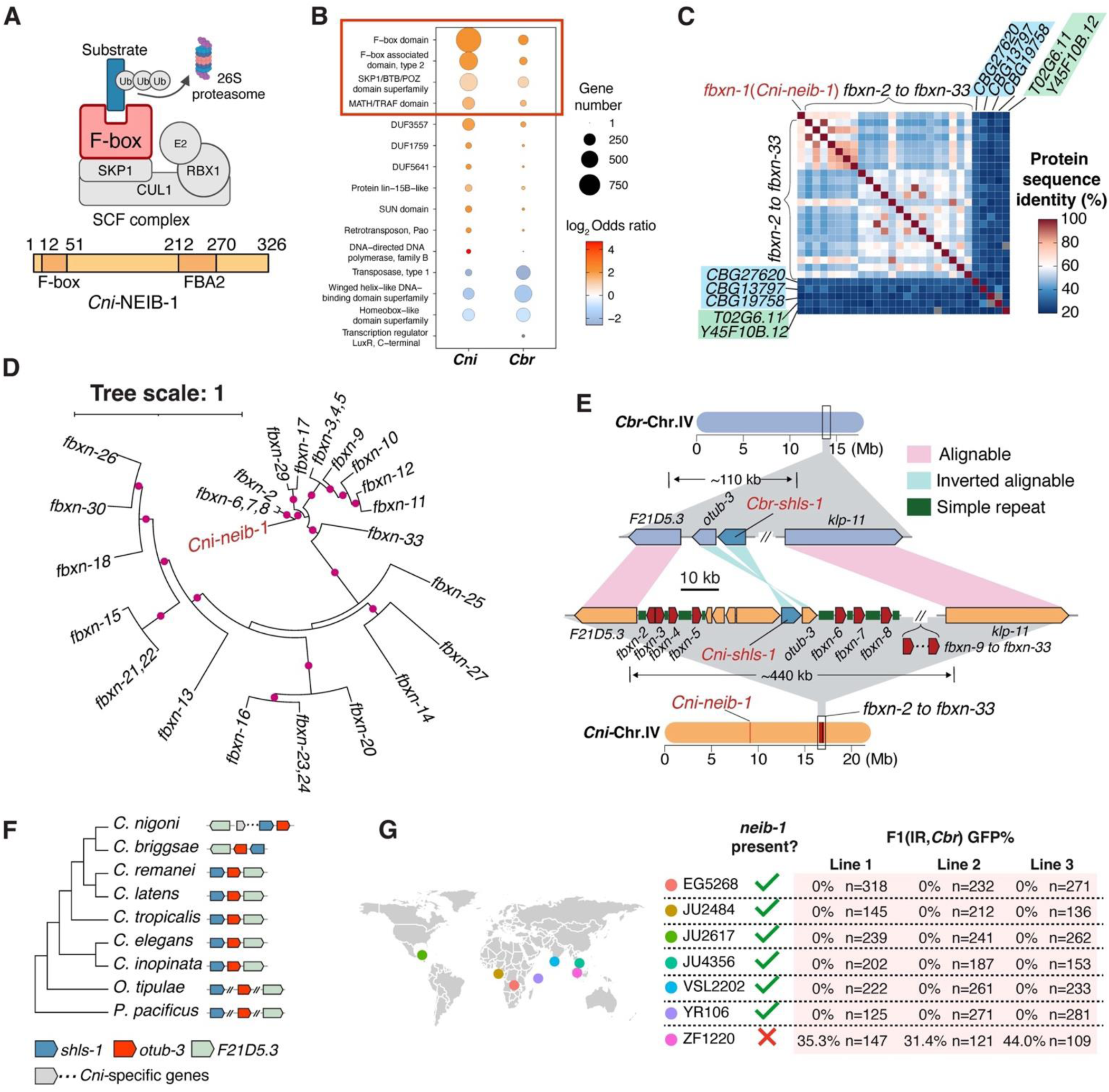
*Cni-neib-1* belongs to a recently expanded, *C. nigoni*-specific F-box gene family. **(A)** Top: Schematic of the ubiquitin-protein ligase (E3 ligase) involving SCF (Skp, Cullin and F-box) complex, with the F-box protein as a substrate adapter. Bottom: Predicted domains within the *Cni-*NEIB-1 protein. **(B)** Bubble plot showing the protein domains encoded by genes with significant number differences between *C. nigoni* and *C. briggsae* (adjusted *P*<0.01, Fisher’s exact test). A Red rectangle highlights the top four expanded domains in *C. nigoni*. **(C)** Heatmap showing protein sequence identity among *fbxn* family genes and their closest homologues in *C. briggsae* and *C. elegans* (shaded in light blue and green, respectively). **(D)** Protein phylogenetic tree of *fbxn* genes. Purple dots denote branches with bootstrap value over 80. **(E)** Comparison of syntenic regions involving *fbxn* genes. The *fbxn* genes are primarily located in a region on the right arm of Chr. IV, characterized by extensive genomic rearrangement, including duplications of *fbxn* gene and an inversion involving *otub-3* and *shls-1*. *Cni-neib-1* was translocated to the center of Chr. IV. *C. elegans* gene names are used for simplicity. **(F)** Orthology and genomic rearrangement of syntenic regions flanking *shls-1* across nematodes. Left: Protein phylogenetic tree of SHLS-1 orthologs in nine nematodes. Right: Syntenic regions involving *shls-1* and two adjacent genes. Note rearrangements in syntenic regions of *Cbr-shls-1* and *Cni-shls-1* compared to other nematodes. **(G)** Variation of *neib-1* alleles among the *C. nigoni* wild isolates. Left: Geographic distribution of sampled *C. nigoni* wild isolates. Right: Comparison of GFP-expression percentages in F1 adults from crosses between *C. briggsae* mothers and *C. nigoni* wild isolate fathers with heterozygous *C. briggsae* Chr. IV right arm introgressions. Three independent introgression lines were generated for each wild isolate. Presence or absence of *neib-1* results in lethal or viable F1 hybrids, indicated by absence or presence of GFP-expressing F1 adults.

A rapid birth-and-death evolution by positive selection frequently causes extensive divergence of F-box genes (27). *fbxn* genes demonstrated an evolutionary pattern consistent with extensive recent duplication and divergence. For example, multiple sequence alignments of *fbxn* proteins revealed various divergent regions including portions of the FBA2 domain (Fig. S15A). By excluding unalignable regions, we identified at least eight sites under positive selection by comparing *Cni-*NEIB-1 with its most closely related paralogues (Fig. S15B). Surprisingly, four groups of F-box genes among the 33 paralogues contain identical cDNA sequences, including *fbxn-3, −4* and *-5*; *fbxn-6, −7* and *-8*; *fbxn-21* and *-23*; and *fbxn-22* and *-24*, indicating fast evolving of *fbxn* genes (Table S2). These identical members are arranged as local tandem duplicates, suggesting they were derived from very recent duplications. Phylogenetic analysis shows that *Cni-neib-1* is most closely related to *fbxn-2* and the three identical paralogs: *fbxn-6, −7,* and *-8* (Fig. 5D).

All the *fbxn* genes are located on the right arm of *C. nigoni* Chr. IV, forming a local gene cluster, except for *Cni-neib-1*, which is near the center of the same chromosome (Fig. 5E). Various repeats were located within a ∼30kb *C. nigoni*-specific region flanking *Cni-neib-*1, including an intact transposon and several pairs of inverted or direct repeats, suggesting remnants of transposition (Fig. S16). Therefore, *Cni-neib-1* might have been captured and translocated to this transposition hotspot. We next examined the genomic region between *Cni-F21D5.3* and *Cni-klp-11*, where all other *fbxn* genes are situated. We observed extensive simple repeats arranged in tandem next to the *fbxn* genes (Fig. S17A), whereas the *C. briggsae* syntenic region (∼110kb) contained fewer repeats (Fig. S17B). The abundant tandem repeats presumably facilitated the duplication of *fbxn* gene through unequal crossover. The ∼440kb region is largely *C. nigoni* specific, except for two orthologous genes (Fig. 5E). Intriguingly, one of these two genes with orthologs in other species is *Cni-shls-1*, apparently underwent an inversion relative to it *C. briggsae* counterpart (Fig. 5E). We compared the syntenic regions containing *shls-1* and its two immediate conserved neighboring genes *otub-3* and *F21D5.3* across nine related nematode species (Fig. 5F). We observed consistent synteny in all *Caenorhabditis* nematodes except in *C. briggsae* and *C. nigoni* (Fig. 5F). This suggests that gene order in other *Caenorhabditis* nematodes represents an ancestral arrangement. An inversion of the three genes appeared to have occurred in the ancestor of *C. briggsae* and *C. nigoni*. *C. nigoni* then presumably underwent another inversion involving *Cni-shls-1* and *Cni-otub-3*, together with the evolution of novel genes. The inversion may have accelerated their species-specific evolution through suppressed recombination, contributing to the specific incompatibility between *Cni-neib-1* and *Cbr-shls-1*.

### *Cni-neib-1* presence is polymorphic among *C. nigoni* populations

Given that *Cni-neib-1* arose from a very recent gene duplication among other paralogs that even consist of multiple members with identical sequences, we wondered whether *Cni-neib-1* also undergoes birth-and-death evolution among *C. nigoni* populations. To this end, we examined the presence or absence of *Cni-neib-1* in seven *C. nigoni* wild isolates collected at different geographic locations (Fig. 5G). Interestingly, we successfully amplified *Cni-neib-1* genomic DNA from six of them but failed to amplify it in one of the isolates, ZF1220, suggesting that the gene either has undergone a very recent loss or has not evolved in ZF1220. To functionally test the absence of *Cni-neib-1* in ZF1220, we examined the hybrid lethality phenotypes between *C. briggsae* and each of the *C. nigoni* wild isolates. Using the same GFP-marker integrated *C. briggsae* as used for the mapping of *Cni-shls-1* and *Cni-neib-1*, we first generated IR strains for each *C. nigoni* intraspecies, each of which carrying a *C. briggsae* Chr. IV right arm introgression fragment, with the syntenic region in the *C. nigoni* genome expected to contain *Cni-shls-1* but not *Cni-neib-1* (Table. S1). We then crossed each of these IR strains with *C. briggsae* mothers. We observed viable hybrid progeny expressing the GFP marker only from IR strains of ZF1220 but not from those of other *C. nigoni* wild isolates (Fig. 5G). These results indicate that *Cni-neib-1* undergoes rapid birth-and-death evolution even among *C. nigoni* populations, which is consequential in causing HI between the two species.

## Discussion

Gene duplications are frequently observed across species and populations, yet their impact on gene flow and the establishment of reproductive barriers remains largely enigmatic (9). The major mechanistic link between gene duplication event and HI was posited to be a reciprocal silencing model involving loss of different copies of duplicated genes (28). Contrary to this assumption, we here identify a novel and *C. nigoni*-specific F-box gene *Cni-neib-1*, derived from a recent duplication, which blocks gene flow from *C. nigoni* to *C. briggsae* via specific destruction of the essential conserved phosphoglucomutase encoded by *shls-1* in *C. briggsae*. Although the presence of *C. nigoni* PGM masks the deleterious interaction, its absence kills the hybrid embryos presumably due to the deficiency of UDP-GlcNAc and deficient glycosylation (Fig. 6A). The incompatibility between *Cni-neib-1* and *Cbr-shls-1* provides evidence of how recently duplicated genes can stop gene flow, thereby either initiating speciation or reenforcing existing speciation barriers. This represents a novel molecular mechanism of DMI model, a concept long proposed with few empirical evidence (4, 5, 29), and demonstrates that HI mechanisms among nematodes extend beyond the toxin-antidote model (22–25).

**Figure 6.**
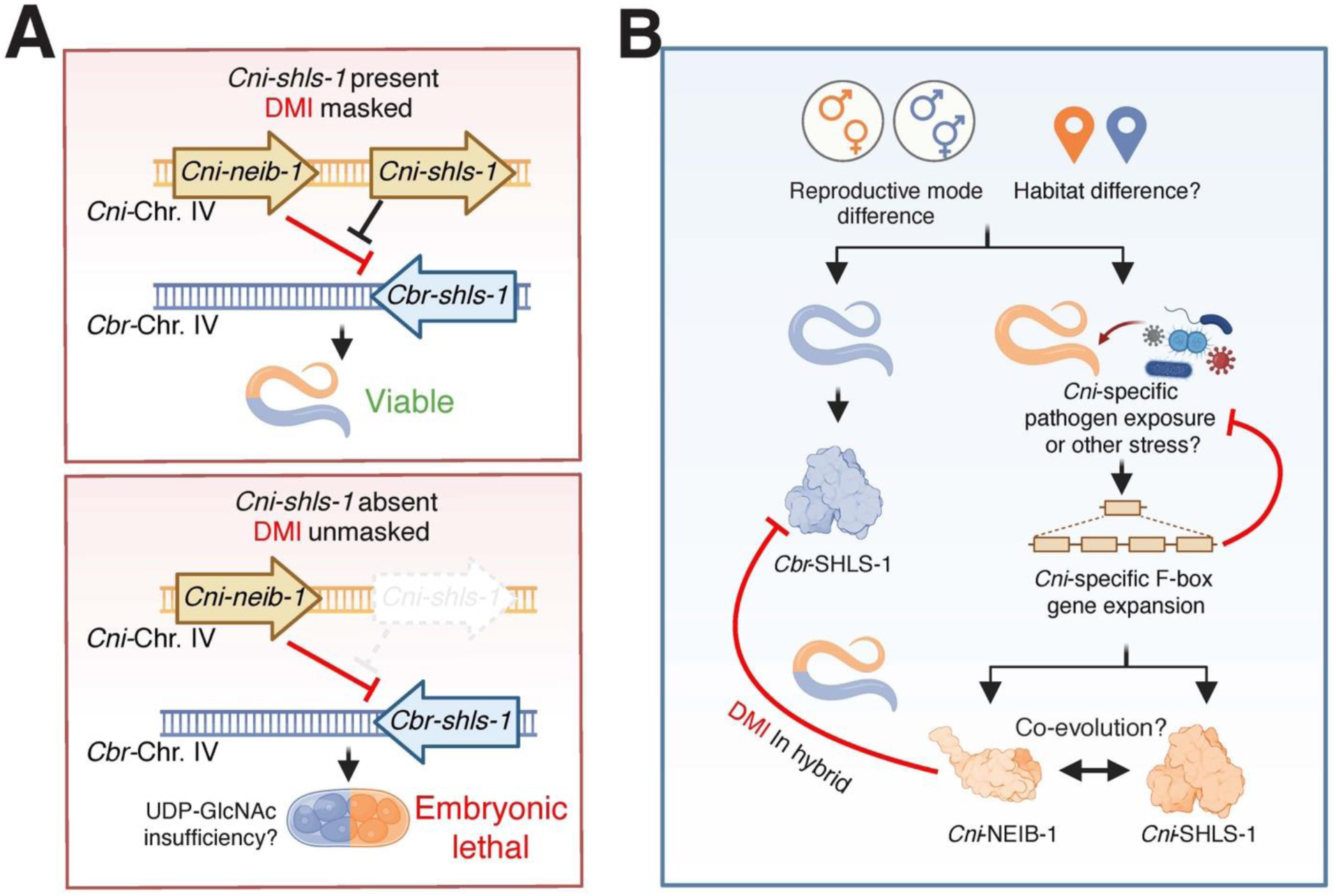
Model of F1 hybrid lethality between *C. nigoni* and *C. briggsae*. **(A)** The interaction between *Cni-neib-1* and *Cbr-shls-1* aligns with the DM model. Top: The presence of *Cni-shls-1* in F1 hybrids masks the negative interaction between *Cni-neib-1* and *Cbr-shls-1*, thereby suppressing the hybrid lethality. Bottom: Without *Cni-shls-1*, the negative interaction depletes *Cbr-*SHLS-1, causing the hybrid embryonic lethality possible through UDP-GlcNAc insufficiency. **(B)** Differences in reproductive mode (gonochoristic in *C. nigoni* and androdiecious in *C. briggsae*) and/or habitat likely expose *C. nigoni* to specific pathogens or stresses, driving species-specific F-box gene expansion, including *Cni-neib-1*, as a defense mechanism. *Cni-neib-1* may accidentally mistargets *Cbr-shls-1* in F1 hybrids, whereas *Cni-shls-1* avoids such mistargeting likely due to co-evolution with *Cni-neib-1*.

Here an essential enzyme may become accidentally degraded by a novel gene product instead of previously reported incompatible components protein complexes (30–32) or epistatic protein-DNA interactions in hybrids (33, 34). In addition, while the canonical DM model posits negative epistatic interactions in evolved alleles of ancestral genes with differential evolutionary history (35), our results demonstrate that the negative epistasis can also occur between a newly evolved gene in one species and a conserved gene in another, and suggest that coevolution may occur between *Cni-neib-1* and *Cni-shls-1* to avoid self-destruction. The involvement of a highly conserved PGM in reproductive isolation also suggests that essential pathways like glucose metabolism are vulnerable to disruptions during reproductive barrier establishment. This is supported by recent findings of incompatible galactose pathway genes in yeast hybrids (36).

F-box proteins are postulated to function as an innate immune response against pathogenic threats, compensating for the lack of adaptive-immunity in nematodes (27). This is supported by a rapid and extensive expansion of the F-box gene family in *Caenorhabditis* nematodes (11, 27). Many F-box genes appear to be under positive selection and exhibit considerable dynamics to fulfill substrate specificity in the E3 ubiquitin system (26). Novel F-box genes have also twice been co-opted during nematode evolution of reproductive mode (37, 38). However, such evolution may inevitably induce unintended protein degradation analogous to an autoimmunity. Therefore, the negative epistatic interaction between *Cni-neib-1* and *Cbr-shls-1* may primarily emerge as a “by-product” of the positive selection of *C. nigoni-*specific F-box genes in arms races with environmental stresses (Fig. 6B). Similar DM incompatibilities involving immune genes have been linked to autoimmune-like hybrid necrosis in plants (39, 40). A key question remains as to why a significant expansion of F-box gene family occurred exclusively in *C. nigoni* but not in *C. briggsae.* One plausible explanation is that the recent evolution of F-box genes, including the *fbxn* family, reflects an adaptive response to species-specific environmental or internal stresses, such as pathogens and transposable elements (Fig. 6B). This may particularly be the case for *C. nigoni*, which requires obligatory mating for reproduction, whereas the hermaphroditic *C. briggsae* is primarily propagated through selfing. Mating behavior can increase the physiological and pathogenic risks in the outcrossing *C. nigoni* (11, 41, 42). Alternatively, the two species may inhabit different niches associated with differential environmental threats. Gene duplications are frequently observed in response to environmental changes across organisms (43), and differential adaptation to environments has also been proposed to trigger a significant expansion of F-box genes in plants (44, 45).

The rapid degradation of *C. briggsae* PGM indicates the high efficiency of *Cni-*NEIB-1, which depletes the enzyme to an undetectable level within 40-60 minutes following the onset of gastrulation in hybrid embryos (Fig. 3). This suggests that zygotic expression of *Cni-neib-1* at the approximately 28-cell stage triggers *Cbr-*SHLS-1 degradation (Fig. 4), aligning with the onset of robust zygotic gene expression during gastrulation (46–49). The specificity of *Cni-* NEIB-1 for *C. briggsae* PGM might stem from some amino acid substitutions, although such sequence changes are rarely documented in HI studies (31, 50, 51). Based on this, co-evolution with *Cni-neib-1* might have led to amino acid substitutions that prevent *Cni-*SHLS-1 degradation by *Cni-*NEIB-1 in *C. nigoni*, a selective pressure absents in *C. briggsae* following reproductive isolation (Fig. 6B). Systematic substituting candidate amino acids could reveal key molecular differences influencing differential targeting by *Cni-*NEIB-1.

Given the pronounced F-box gene expansion in a species-specific pattern, we finally propose a “creation by degradation” model. That is, selective degradation of specific targets could drive evolutionary novelties, enhancing survival against environmental stresses, competing pathogens or optimizing reproductive strategies, thus facilitating speciation. This is particularly relevant to the immune responses that are prevalent across all organisms, for example, via genetic changes in microorganisms (52) and adaptive immunity involving somatic mutation through antibodies in vertebrates (53), both of which can be associated with extensive gene duplications. Selective degradation of foreign elements imposes significant selection pressures on hosts, leading to a balance between fighting against “non-self” and accidentally targeting “self” (54). We anticipate that degradation pathway proteins, such as F-box proteins, are pivotal in such evolutionary processes.

## Materials and Methods

### Nematode maintenance and reference strains for *C. briggsae* and *C. nigoni*

All worms were maintained at 25 °C on modified culture plates with a two-fold increase in agar concentration of standard nematode growth medium (NGM) plates, to prevent *C. briggsae* from burrowing into the agar. JU1421 was used as the *C. nigoni* wild-type (WT) reference strain (14, 17, 55) for the generation of IR strains and conducting interspecies crossing, except for those involving *C. nigoni* wild isolates. *C. briggsae* (AF16) reproduces mostly by selfing of hermaphrodites that produce their own oocytes and sperm. It is therefore challenging to distinguish between its selfing progeny and F1 hybrids with *C. nigoni* fathers, complicating the phenotyping of F1 hybrids. To circumvent this, a homozygous mutant strain, *Cbr-she-1* (*V49*) (38), in an AF16 background was used as the WT *C. briggsae* reference strain for all crosses, phenotyping, genome editing and expression analysis. The *Cbr-she-1* mutant exhibits no apparent defects relative to AF16 strain, except for a feminized germline that produced no sperm (38), thereby effectively functioning as a *C. briggsae* strain with both males and females.

### Generation of introgression (IR) strains

All the IR strains were generated by repeatedly backcrossing individual transgenic *C. briggsae*, which was labeled with a chromosomally integrated fluorescent marker, into *C. nigoni* for at least 15 generations as previously detailed (14) and in Fig. 1B. The GFP or mCherry marker (*pmyo-2::gfp* and *pmyo-2::mCherry*, pharynx-expressed) was inserted to a random location on one of the six *C. briggsae* chromosomes using either biolistic bombardment or *miniMos* transgenesis method (14, 56) (Fig. 1B). We generated over 100 independent *C. briggsae* transgenic lines, each carrying a single marker insertion. Due to spontaneous recombination and random chromosome segregation (Fig. 1B), each repeated backcross resulted in an IR strain with a single *C. briggsae* fragment linked to the fluorescent marker as a heterozygote in an otherwise *C. nigoni* background. The GFP transgenic marker in ZZY10018, integrated using the biolistic bombardment method, was inserted on the right arm of Chr. IV, with its integration site roughly mapped to ∼13.5Mb (32). All the other IR strains carrying the *C. briggsae* Chr. IV introgression were generated by repeatedly backcrossing the *C. briggsae* transgenic strain ZZY0734, which carries a GFP marker on the right arm of Chr. IV, to *C. nigoni*. The strain ZZY10413 carries a single-copy GFP marker, integrated at 16.98 Mb on *C. briggsae* Chr. IV using the *miniMos* method (57). The generation of *C. briggsae* introgressions with precise recombination breakpoints for mapping of *Cni-shls-1* and *Cni-neib-1* was achieved using CRISPR/Cas9-mediated targeted recombination (see “Gene mapping and cloning through targeted recombination with CRISPR/Cas9”). All IR strains with dual *C. briggsae* introgressions (Fig. 4D and E) were generated by crossing the GFP-labeled IR strain ZZY10413 (Fig. 2A), with another IR strain labeled with mCherry, integrated at various locations of *C. briggsae* genome (Fig. S10). ZZY10423 was generated using a *C. briggsae* strain labeled with both GFP and mCherry markers, resulting from the cross between ZZY0734 and ZZY0765 (mCherry marker integrated at ∼5.93Mb on *C. briggsae* Chr. IV). The mCherry markers was also integrated in the *C. briggsae* genome using the *miniMos* method, and all the mCherry-labeled IR strains were generated through spontaneous recombination. The IR strains for various *C. nigoni* wild isolates were generated by repeatedly backcrossing ZZY0734 with each wild isolate.

For IR strains not generated by targeted recombination, the approximate size of the introgression was determined by assessing the presence or absence of *C. briggsae* genomic DNA using single worm PCR (swPCR) to amplify genomic fragments, as previously described (14). All the genomic coordinates in this study were based on our newly assembled genome of *C. nigoni* (CN3) and *C. briggsae* (CB5) (18). A comprehensive list of IR strains used in this study, along with their starting *C. briggsae* transgenic strains, genotypes, F1 hybrid phenotypes (F1 hybrid lethality or not indicated by the presence or absence of GFP-expressing hybrid adults), and introgression coordinates, is detailed in Table S1. All swPCR primers used for genetic mapping and determining introgression sizes in this study, except those previously reported (14, 17), are listed in Table S3.

### Phenotyping F1 embryonic lethality using IR strains

Direct measurement of hybrid embryo lethality of progeny between *C. briggsae* and an IR strain carrying a single or dual heterozygous introgression fragment can be tedious, because it produces two cohorts of F1 hybrids carrying two different genotypes: WT and introgression-bearing (Fluorescence-expressing F1 hybrids with *C. briggsae* introgression fragment as homozygotes) ones (Fig. 1B and D; Fig. 4C). Nearly 50% of the WT F1 hybrids were also arrested as embryos from reciprocal crosses (Fig. S1A), which complicates the measurement of embryonic lethality percentages. We therefore used the absence of GFP-expressing hybrid F1 adults as a proxy for the lethality of GFP-expressing hybrid embryos (Fig. 1C and D).

The IR strains we generated previously carry various introgression fragments from different parts of the *C. briggsae* genome, cumulatively covering all the *C. briggsae* genome (14). Therefore, crosses between these IR strains with *C. briggsae* produced hybrid F1 progeny that, cumulatively, lacked the entire *C. nigoni* genome, allowing for the initial genome-wide screening of hybrid lethality suppressor loci. Similarly, because the dual introgression bearing IR strains carry either the left and the right half of a *C. briggsae* autosome other than Chr. IV, and the two introgressions were overlapping with each other (Fig. S10), the cumulative introgression fragments were expected to cover all *C. briggsae* autosomes except for Chr. IV. Therefore, the cumulative absence of syntenic *C. nigoni* genome regions covers the entire *C. nigoni* genome other than Chr. IV, allowing for a genome-wide screening of *Cni-neib-1*.

### Crosses, phenotyping and statistical analysis of the crossing results

To characterize the hybrid embryonic lethality phenotype, each cross was performed as follows: a minimum of fifty L4-stage females and twenty young adult males were picked onto a 55-mm modified NGM plate. OP50 bacteria were seeded only in the center of the plate to facilitate overnight mating. Each cross was performed at least twice. On the following day, fertilized females were transferred to at least three 35-mm modified NGM plates, each similarly seeded with OP50 in the center and containing ten animals and allowed to lay eggs for approximately 3 hours. The females were then removed. All laid embryos were counted to determine the total number of eggs (n). The number of unhatched embryos in each plate was counted again after around 24 hours to assess the hatching rate. The number of adult worms, with or without GFP expression, was counted two days later using a Leica M165FC fluorescent stereo microscope, when all the hatched animals were expected to reach adulthood. The presence of arrested larvae was also counted under a regular stereo microscope when all the worms were expected to reach adulthood. For all the crosses involving animals carrying an extrachromosomal array, progeny that inherited the array were identified by the observation of the injection marker expression (pZZ184, see “Generation of extrachromosomal array lines”). The 95% confidence intervals of the ratios of hatching rate and larval arrest percentage were calculated using the Agresti-Coull method and represented by the error bars. The significance of the difference in ratios (*P*-value) was evaluated using either the chi-square test or the Fisher’s exact test, where applicable.

### Gene mapping and cloning through targeted recombination with CRISPR/Cas9

We previously demonstrated extensive recombination suppression between the two species, which prevents an efficient generation of IR strains with desirable introgression sizes (12, 18). To facilitate the mapping and cloning of *Cni-shls-1* and *Cni-neib-1*, targeted recombination was performed using single or dual gRNAs to induce targeted double-strand DNA breaks and subsequent crossovers, as previously described (18). Single gRNA was applied to trigger recombination in the initial F1 hybrids from the introgression processes, and successful recombination at the target sites was maintained during each round of repeated backcrossing to generate the final IR strain. Dual gRNAs were used on an IR strain with a relatively large *C. briggsae* introgression fragment to generate IR strain with a desirable introgression length, as illustrated in Fig. 1E. Targeted recombinants were screened by amplifying *C. briggsae* genomic fragments flanking the expected recombination breakpoint using swPCR, as previously described (18). The recombination site was always re-verified by swPCR before each interspecies crossing to prevent any further spontaneous recombination during worm maintenance. Further recombination was extremely rare in all Chr. IV IR strains, consistent with our previous observations that recombination predominantly occurred in the F1 or F2 generations during the generation of IR strains (12). All the introgressions in the IR strains for fine-mapping of *Cni-shls-1* and *Cni-neib-1* were extended to the right-end of *C. briggsae* Chr. IV, as confirmed by swPCR, due to the insertion of GFP marker at the distal right end of *C. briggsae* Chr. IV. Consequently, IR strains with a larger introgression fragment of *C. briggsae* Chr. IV possessed a smaller *C. nigoni* genomic fragment on the introgression-bearing chromosome, and vice versa. The candidate interval was deduced by excluding the overlapping regions of *C. nigoni* Chr. IV on the introgression-bearing chromosomes in two strains with differential hybrid phenotypes, i.e., the presence or absence of GFP-expressing hybrid adults (Fig. 1E). All gRNA sequences used for targeted recombination, along with the resulting strains, are listed in Table S4.

### Generation of extrachromosomal array lines

The overexpression constructs for *Cni-shls-1*, pZZ220 [*Peft-3::Cni-shls-1 CDS::tbb-2 3’UTR*; NeoR], and *Cbr-shls-1*, pZZ233 [*Peft-3::Cbr-shls-1 CDS::tbb-2 3’UTR*; NeoR], were generated by replacing the GFP coding sequence in pCFJ914 with the *Cni-shls-1* or *Cbr-shls-1* coding sequences, respectively. Specifically, total RNA from *C. briggsae* and *C. nigoni* mixed stage worms was extracted using the TRIzol (Invitrogen) method, following the manufacturer’s instructions. cDNA was reverse transcribed using the PrimeScript RT Reagent Kit (Takara). The coding sequences of *Cni-shls-1* and *Cbr-shls-1* were then amplified, using the cDNA library as the templates with the following primers: *Cni-shls-*1 Forward: 5’-ATGACAAATCTCGATTTTCCACCCCAATACACCCG-3’; Reverse: 5’-CTAAGCAGAACTAAGGCTCAGGACGACTTGTTCCA-3’; *Cbr-shls-*1 Forward: 5’-ATGGCGAATCTCGAATTTCCACCCCAATACACC-3’; Reverse: 5’-CTAAGCAGAACCAATGCTGAGGACGACTTGTTC-3’. The PCR-amplified products were treated with T4 Polynucleotide Kinase (NEB) for phosphorylation and then blunt-end ligated to the backbone amplified from pCFJ914, excluding the GFP sequence. The correct orientation was validated by PCR amplification of the insertion boundary followed by Sanger sequencing. The transgene for overexpression of *Cni-neib-1* was constructed using its endogenous genomic sequence. The genomic sequence of *Cni-neib-1* was amplified using the following primers: Forward: 5’-TGATACTGTACCTTGCTCAGTAGAC-3’; Reverse: 5’-CAGAAGTATCCATTGTCGGGAGATC-3’. The same primer pair was used to amplify the genomic sequence containing *Cni-neib-1* in *C. nigoni* wild isolates. The injection mixture was prepared by mixing DNA constructs or PCR products with the injection marker at a 3:1 ratio, achieving a final concentration of 40 ng/µl. Both sides of the gonad were injected following the standard worm microinjection protocol. To generate all the extrachromosomal array lines, pZZ184 (57) was used as the injection marker, driving mCherry expression in the pharyngeal muscle of worms. The sequence of pZZ184, pZZ220 and pZZ233 can be found on Benchling (pZZ184: https://benchling.com/s/seq-7mm2RrWOlhXQekND4YsV?m=slm-afbR5xmfnS4uJZuboWap; pZZ220: https://benchling.com/s/seq-rLFZKakJP9pUMKuvk6qW?m=slm-JGdUaCYJwLeNtnaDENWO; pZZ233: https://benchling.com/s/seq-Rw1FSJqtM6tFUa1vAfyu?m=slm-Yig0xmvXNYpV6Vp24knx).

### Genome editing for the generation of knockout and knock-in strains using CRISPR/Cas9

All the CRISPR/Cas9 mediated genome editing was performed as follows: All tracrRNA and crRNA were acquired from IDT and dissolved in the duplex buffer (IDT) to a final concentration of 200 µM. gRNA duplexes were prepared by mixing 2 µL each of tracrRNA and crRNA, followed by incubation at 95°C for 5 minutes and gradual cooling to room temperature in a thermal cycler (Life Sciences) following the melting DNA method (58). Each 2 µL of the gRNA duplex was thoroughly mixed with an equal volume of Cas9 proteins (IDT, glycerol-free, 10 µg/µL) and 2 µL of the injection marker (30 ng/µL, dissolved in nuclease-free water). The mixture was then incubated on ice to allow the formation of the ribonucleoprotein (RNP) complex before being loaded into the microinjection needles. For editing with dual gRNAs, RNP complexes were prepared similarly, except that 1 µL of each gRNA duplex was added. Deletion of large genomic fragments encompassing multiple genes, or smaller fragments containing a single gene, was routinely achieved using dual gRNAs. To generate the *Cni-shls-1* and *Cni-neib-1* double mutant strains, the *Cni-neib-1* homozygous mutants (ZZY1073 and ZZY1095) were first generated. *Cni-shls-1* was subsequently knocked out in the ZZY1073 and ZZY1095 backgrounds respectively, resulting in the double knockout strains ZZY1096 and ZZY1097. Successful deletions were verified by swPCR amplification of the deletion boundary, which was not expected to be amplified in worms without successful editing. To validate the deletion, amplified fragments spanning the deletion boundaries were purified using FastPure Gel DNA Extraction Mini Kit (Vazyme) before being sent for Sanger sequencing, with the amplifying primers used as sequencing primers. The deletion boundaries were visualized on Benchling (https://www.benchling.com/). All the deletion alleles generated for the molecular cloning and characterization of *Cni-shls-1* and *Cni-neib-1*, verified by Sanger sequencing are shown in Fig. S18 and Fig. S19, respectively. To knock in fluorescent markers at the N terminus of *Cni-shls-1* and *Cbr-shls-1*, the coding sequence for *gfp* and *mCherry*, along with the 3’ linker sequences, were amplified from pZZ160 and pZZ184 (pZZ160: https://benchling.com/s/seq-chx4OG5kSsm1rxOikYhl?m=slm-8qY94UE9qNe019pA7GcZ), respectively. A 30-nt homologous arm was added to the 5’ end of each primer, and the ends were modified with a 5’ Spacer 9 (SP9) modification, as recommended by Ghanta and Mello (58). The DNA amplicon was subjected to a melting process as described by Ghanta et al. to improve knock-in efficiency (58). For substitution of the divergent motif sequence in *C. briggsae shls-1* with that from *C. nigoni*, a repair template, including the *C. nigoni* motif sequence along with 30-nt *C. briggsae* homologous arms, was synthesized as single-stranded DNA from IDT. The melted PCR products or donor oligos were mixed with Cas9 RNP complex to a final concentration of 200 µM. The gRNAs used for the generation of deletion or knock-in mutants, the primers used for amplifying the deletion boundaries or the successful knock-in boundaries, and the genome-edited strains generated in this study were listed in Table S4, S5 and S6, respectively.

### Generation of knock-in strain with *MiniMos*

To knock in *Cni-shls-1* into *C. briggsae*, the *Cni-shls-1* transgene construct pZZ220 (see “Generation of extrachromosomal array lines”) was chromosomally integrated into the AF16 strain using the *miniMos* method. The insertion sites were determined using inverse PCR as described (59). Two independent transgenic strains, i.e., ZZY1005 and ZZY1006, were generated with insertion sites located at positions 10,285,962 on Chr. I and 4,563,560 on Chr. V, respectively. The two transgenes were crossed into the *Cbr-she-1* (*V49*) mutant background to generate strains ZZY1009 and ZZY1010, respectively.

### Testing the incompatible effects of *Cni-neib-1* DNA in *C. briggsae* and *C. nigoni*

*Cni-neib-1* genomic DNA was amplified using the same primer pairs for generating the *Cni-neib-1* array (see “Generation of extrachromosomal array lines”). The PCR amplicon was purified with using FastPure Gel DNA Extraction Mini Kit (Vazyme) and dissolved in nuclease-free water to a final concentration of 20 ng/µl before being injected into 15 to 20 *C. briggsae*, *C. briggsae* with a *Cni-shls-1* knock-in and *C. nigoni*, which were then subjected to recovery for three hours. The embryos laid over six hours were characterized for the proportion of hatching. As a control, nuclease-free water was injected in *C. briggsae* and *C. nigoni*, and the hatching rates of their progeny were quantified.

### RNA interference

Genomic DNAs of mixed-stage worms was extracted using the PureLink Genomic DNA Mini Kit (Invitrogen). Partial genomic sequences of *Cni-shls-1* and *Cbr-shls-1* were PCR-amplified using primers flanked by T7 promoter sequences, i.e., forward: 5’-taatacgactcactatagggTGCCGTTCATCGTCTATCGG-3’ and reverse: 5’-taatacgactcactatagggAACCTTGATGCGTTGCTTCG-3’ for *Cni-shls-1*; forward: 5’-taatacgactcactatagggTGCCGGCCAACTTCATTTTG-3’ and reverse: 5’-taatacgactcactatagggTCTGCGAACGACGGAATCAA-3’ for *Cbr-shls-1*. Double-stranded RNAs (dsRNAs) were *in vitro* transcribed using the HiScribe T7 Quick High Yield RNA Synthesis kit (NEB) as described (60). The dsRNAs were diluted to a concentration of 100 ng/µl with nuclease-free water and injected into both gonads of fertilized young adult females that had been mated with males of their own species. Six to eight injected females were placed onto a 55-mm crossing plate and transferred to a new plate at 12-hour intervals. The ratios of hatching embryos and arrested larval were calculated for the first two consecutive 12 periods, as all the injected females became sterile after 24 hours. The number of arrested larvae was determined by subtracting the number of adults from the number of hatched eggs. The larval arrest ratio was calculated accordingly.

### Native-RNA sequencing and data analyses

*C. nigoni* embryos were collected by bleaching gravid worms on two 90-mm culture plates densely populated with adults. The embryos were washed at least three times with M9 buffer before TRIzol reagent (Invitrogen) was added. To ensure the complete lysis, the embryos underwent at least 10 freeze-and-thaw cycles. Total RNA was then extracted following the manufacturer’s instruction. Each 5 µg of extracted total RNA was used for library preparation using the SQK-RNA002 kit (Oxford Nanopore Technologies, ONT). The prepared library was loaded onto a FLO-MIN106D flow cell and sequenced on an in-house MinION device (ONT), following the standard MinKNOW (v19.12.5) protocol for at least 48 hours. Raw sequencing signal files (FAST5) were basecalled by Guppy (v4.0.11) using the high-accuracy model. The “–qscore_filtering” option was applied to categorize reads as “pass” or “fail” with the default cutoff. Only passed reads were retained for further analysis. Illumina RNA-seq data for *C. nigoni* embryos were retrieved from our previously published dataset (15). The reads were mapped to *C. nigoni* genome (CN3) using minimap2 (v2. 17) (61) for the ONT reads and Star (v2.7.0b) (62) for the Illumina RNA-seq reads. Read alignments were extracted using SAMtools (v1.9) (62) and visualized with the Integrative Genomics Viewer (IGV, v2.16.0) (63).

### Gene annotation and the identification of *fbxn* genes

The repeat files for *C. briggsae* and *C. nigoni* were generated using RepeatModeler (v2.0.3) with default parameters. We found that most duplicated genes were categorized as repeats. To prevent the masking of these gene and maximize the protein-coding gene prediction, we excluded all repeat families containing protein-coding genes for gene prediction similarly as others did previously (11). Specifically, we predicted all possible open reading frames within the repeats using “getorf” from EMBOSS (v6.6.0) with the argument “-minsize 90” and extracted those encoding more than 30 amino acids. We then aligned all these peptides to the *C. elegans* proteome from Wormbase (WS280) (64) using BLASTP (v2.16.0) with the parameters “-evalue 1e-09 -seg yes”. The domains within the peptides were also predicted using InterProScan (v5.65-97). Repeats containing peptides that aligned to any member of the *C. elegans* proteome or had predicted protein domains were removed from the repeat files. The remaining repeats were used as input for genome masking with RepeatMasker (4.1.2) using the argument “-e ncbi -s xsmall -gccalc -gff”. Gene models were predicted using AUGUSTUS (v3.4.0) (65) on the CN3 genome with the arguments “--gff3=on --progress=true -- species=caenorhabditis --uniqueGeneId=true --protein=on --introns=on --start=on --stop=on - -cds=on --codingseq=on Genome.fa”. Coding DNA and protein sequences were extracted using the getAnnoFasta.pl script from AUGUSTUS. The *fbxn* gene paralogs were identified by OrthoFinder (v2.5.4) (66) using default parameters. All predicted *fbxn* gene models were verified and curated manually by analyzing both ONT and NGS RNA-seq reads. Genes were categorized as potential pseudogenes if their structure had undergone remarkable changes, such as abnormally large introns or local duplications of exons and introns. All the *fbxn* genes are listed in Table S2.

### Gene domain prediction and orthogroup classification

The longest isoform of all annotated genes was extracted using the agat_sp_keep_longest_isoform.pl script from the AGAT toolkit (v1.0.0) and used as the input for InterProScan to predict protein domains. The number of genes associated with each predicted protein domain was calculated. Differences in gene number between *C. briggsae* and *C. nigoni* for each protein domain were compared using Fisher’s exact test, with adjusted *P*<0.01 (using FDR multiple test correction) considered as significant. The log2 odds ratio for each protein family was also calculated for the generation of the bubble plot (Fig. 5B).

### Sequence alignment and phylogenetic analyses

Pairwise alignment between the protein sequences of *Cni-shls-1* and *Cbr-shls-1* was performed using the EMBOSS Needle pairwise alignment tool (67). The active sites of SHLS-1 were predicted using NCBI Conserved Domain Search (68). The schematics of the alignment results shown in Fig. 3A was plotted using ggplot2 in R. Multiple sequence alignments of nematode SHLS-1 and the F-box proteins were performed using Clustal Omega (69), and the results were visualized using Jalview (v2.11.3.2) (70). The sequences of SHLS-1 orthologs were retrieved from WormBase (WS280) (64). Nine nematodes were selected for SHLS-1 sequence alignment because their genomes were accurately assembled to the chromosome level using long-read sequencing technology. Phylogenetic trees of *fbxn* proteins and nematode SHLS-1 were constructed using IQ-TREE (71) with the maximum-likelihood method using the command “-m MFP -bb 1000 -alrt 1000”. The tree results were visualized and adapted from iTOL (72). The *d_N_/d_S_* ratio of *Cni-*NEIB-1 and its closest *fbxn* proteins was calculated using EasyCodeML (73) with model 8 of the CodeML program from PAML (74). All DNA sequence alignments were performed using BLASTn (75) with the argument “-word_size 7” to include all short alignments.

### Protein structure prediction and structural alignment

The protein structures of *C. briggsae* and *C. nigoni* SHLS-1, as well as human PGM3, were predicted by AlphaFold 2 via ColabFold (v1.5.5) (76) and visualized using PyMOL (v2.5.5). Structural alignments were performed using the align function in PyMOL, and pairwise alignment RMSD (root mean square deviation) values were retrieved.

### Imaging

Low-resolution micrographs of ZZY10018 adults and its hybrids with *C. briggsae* were taken using an M165FC Fluorescent Stereo Microscope (Leica). High-resolution DIC and fluorescent images were acquired using a Stellaris confocal microscope (Leica). A 60x water immersion objective was used, except for the hybrid embryos from ZZY10018, for which a 20x objective was applied. To acquire images of RNAi-treated *C. nigoni* and *C. briggsae* females as well as GFP::*Cni-*SHLS-1 and mCherry::*Cbr-*SHLS-1 males and females, approximately one-day-old adult worms were picked and paralyzed using 10mM sodium azide. Whole worm images were acquired using the tile scanning function. For imaging embryos, 20-30 gravid females were dissected, and the released embryos were transferred to a slide with a pre-dried polylysine (0.5mg/ml) pad at the center to prevent movement during imaging. GFP and mCherry images were acquired using 488 nm and 561 nm laser, respectively, with 3% power and a gain value of 300. All the images were taken (frame size: 712×512 pixels) with a scanning speed of 100Hz and a pinhole size of 2.5. To examine the fluorescence intensity dynamics over development, each fluorescent marker-labeled embryo was imaged at four different development stages: 4-cells stage, gastrulation onset, 40-60 minutes after gastrulation onset and comma stage.

For the time-series imaging of embryogenesis, 2% laser power was applied for both GFP and RFP channels. To better visualize the degradation of *Cbr-*SHLS-1 protein, four different imaging blocks with varying cycles and pause intervals were applied for each embryo: Block 1: 5 cycles, 4-minute pause; Block 2: 30 cycles, 1-minute pause; Block 3: 10 cycles, 4-minute pause and Block 4: 30 cycles, 20-minute pause. The fluorescent intensity of GFP::*Cni-*SHLS-1 and mCherry::*Cbr-*SHLS-1 embryos were measured using ImageJ (v1.8.0). A Corrected Total Cell Fluorescence method was applied to each measurement to remove background signals and correct for intensity differences across areas.

### *C. nigoni* isolate collection and screening

*Caenorhabditis* nematodes were isolated from decomposing vegetal matter as described in (77, 78). Species identification of *C. nigoni* was performed using rDNA ITS2 sequencing as in (78) and test crosses to *C. nigoni* JU1325 or the derived inbred strain JU1422.

### Plots and schematic graph drawing

All the bar plots and line plots were created using GraphPad Prism 10. The candidate intervals, gene models, and their corresponding read tracks and coverage were visualized and adapted from IGV. All schematics were first generated with BioRender (bioRender.com) and further modified using Adobe Illustrator. Dot plots were constructed using ggplot2 in R, with RStudio (v2023.06.0) (79).

## Data and code availability

The genome assembly and annotation files for CN3 and CB5 are accessible in the NCBI Bioprojects database under the accession number PRJNA917437 and on GitHub (https://github.com/PikaPatch/CB-CN). The *C. nigoni* ONT native RNA-seq raw reads are available under the accession number PRJNA1068783. All the raw data and custom scripts used in generating the results are available on Zenodo at https://zenodo.org/records/13299583.

## Acknowledgements

We thank E. Haag and R. Ellis for the *C. briggsae she-1* mutant strain; and Luc Barre, Lise Frézal, Mirko Francesconi, Remy Froissart, Takao Inoue and Varsha Singh for helping in collecting *C. nigoni* wild isolates. We thank Dr. Cindy Tan and Mr. Chung Wai Shing for logistic support and members of Z.Z.’s laboratory for constructive comments.

## Supplementary Materials

Figs. S1 to S19

Tables S1 to S6

Movie S1 to S8

## The PDF file includes

Figs. S1 to S19

Tables S2, S4 to S6

Legends for Movies S1 to S8

## Other Supplementary Materials for this manuscript include the following

Tables S1 and S3

Movie S1 to S8

---

**Figure S1.**
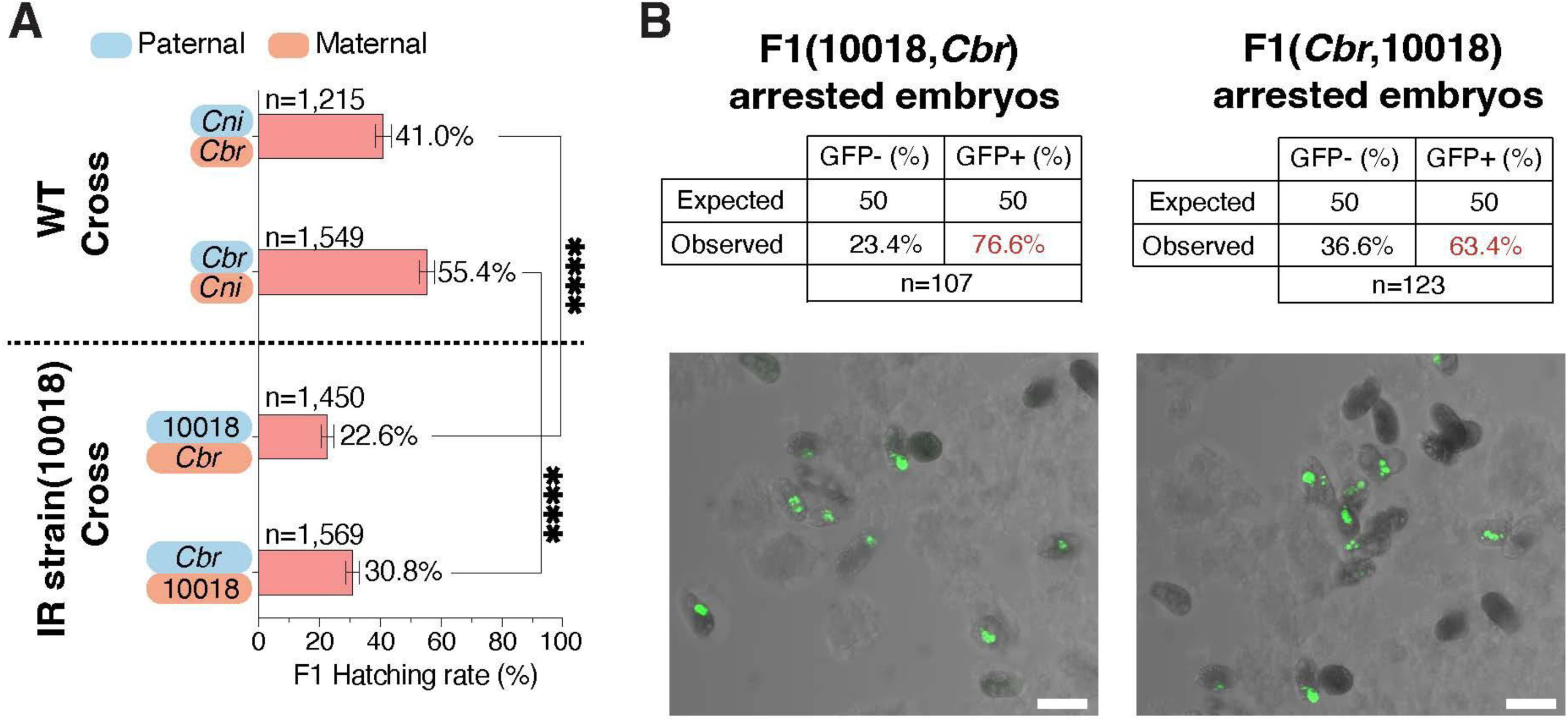
Absence of *C. nigoni* Chr. IV right arm results in embryonic lethality in F1 hybrids. **(A)** Comparison of hatching rates for F1 hybrids from reciprocal crosses of WT parents and *C. briggsae* with IR strain, ZZY10018 (see Fig. 1D), which carries a heterozygous introgression fragment from *C. briggsae* Chr. IV right arm. Note that the hatching rates decrease by approximately 50% decrease in ZZY10018 crosses compared to the WT crosses, regardless of crossing directions. \*\*\*\**P*<0.0001, Fisher’s exact test; error bars: 95% confidence. **(B)** Most arrested F1 hybrid embryos express GFP. Top: Expected and observed proportion of arrested hybrid F1 embryos carrying (GFP+) or lacking (GFP-) GFP marker from ZZY10018 fathers (left) or mothers (right). Bottom: Representative images of arrested hybrid F1 embryos from ZZY10018 fathers (left) or mothers (right). Scale bar: 30µm

**Figure S2.**
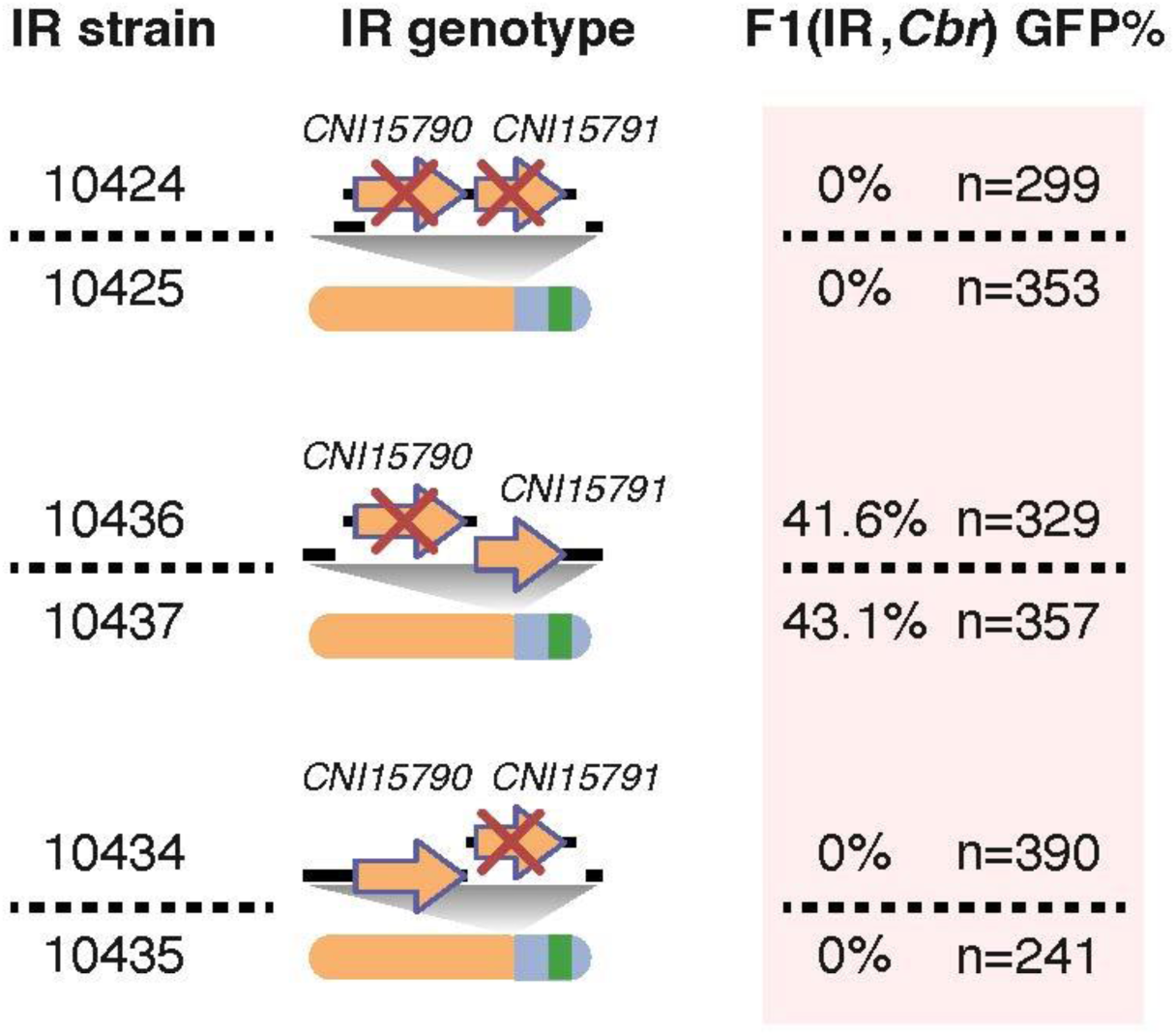
Cloning of *Cni-shls-1* gene through deletion(s) of candidate gene(s) in the ∼33kb mapped interval (see Fig. 2A) Shown are the percentages of GFP-expressing hybrid F1 adults from *C. briggsae* mothers and IR strain fathers with genomic deletion(s) of both genes (top), *CNI15790* only (middle), or *CNI15791* only (bottom) on *C. nigoni* Chr. IV carrying the *C. briggsae* introgression. The deletions are generated in IR strain ZZY10413 (see Fig. 2A). Genotypes schematics for *Cni-*Chr. IVs with the *C. briggsae* introgressions are shown. Two independent lines were generated for each deletion.

**Figure S3.**
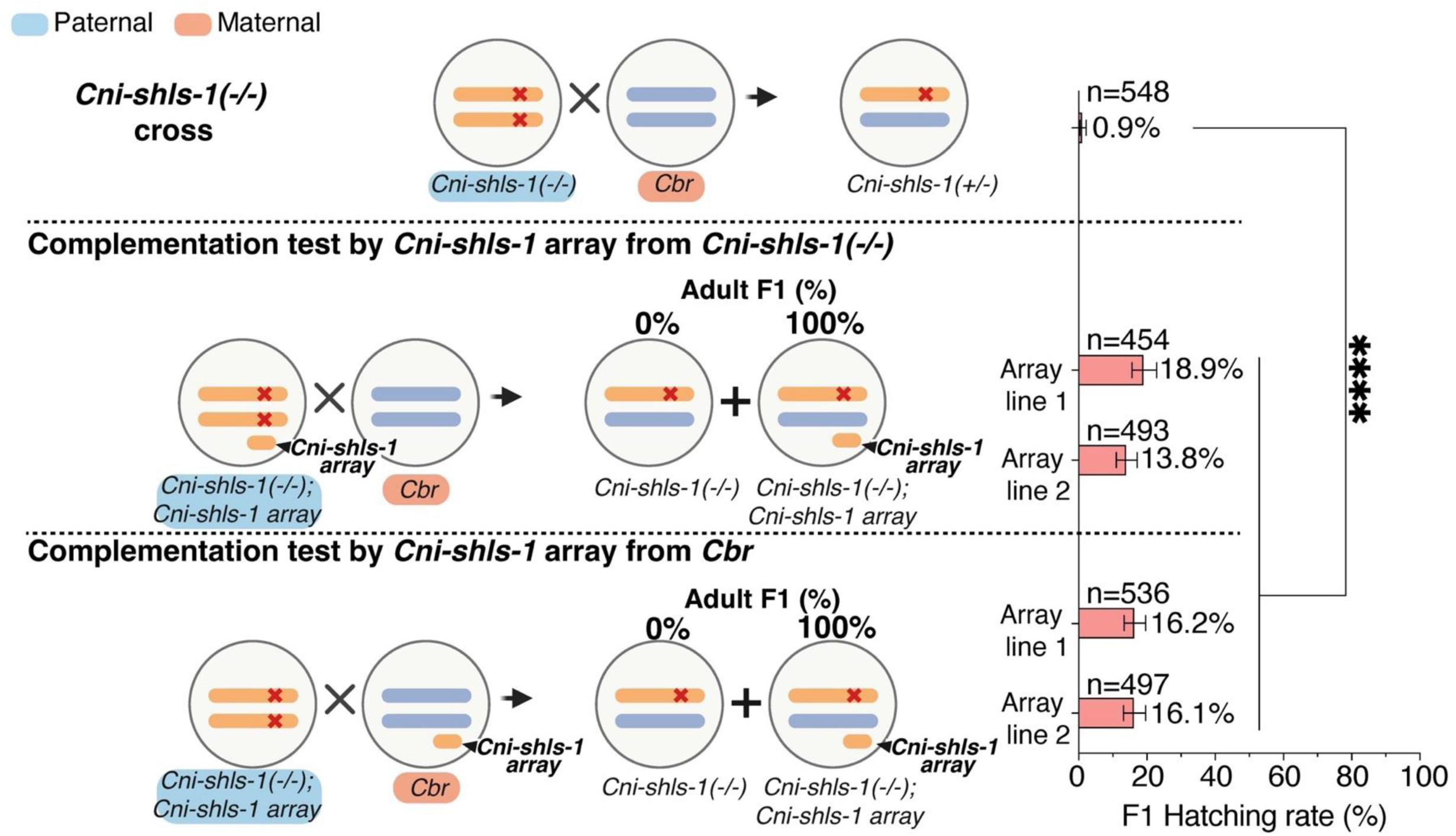
Validation of *Cni-shls-1* as a hybrid lethality suppressor gene. Shown is the comparison of hatching rates for F1 hybrids from *Cni-shls-1(-/-)* fathers and *C. briggsae* mothers (top), and F1 hybrids carrying a *Cni-shls-1* extrachromosomal array inherited from either the *Cni-shls-1(-/-)* fathers (middle) or the *C. briggsae* mothers (bottom). Genotype Schematic (Chr. IV only) for parents and F1 hybrids are shown on the left. Also shown are the percentages of F1 hybrids with/without the array. Two independent array-bearing lines were used for each cross. \*\*\*\**P*<0.0001, Fisher’s exact test; error bars: 95% confidence interval.

**Figure S4.**
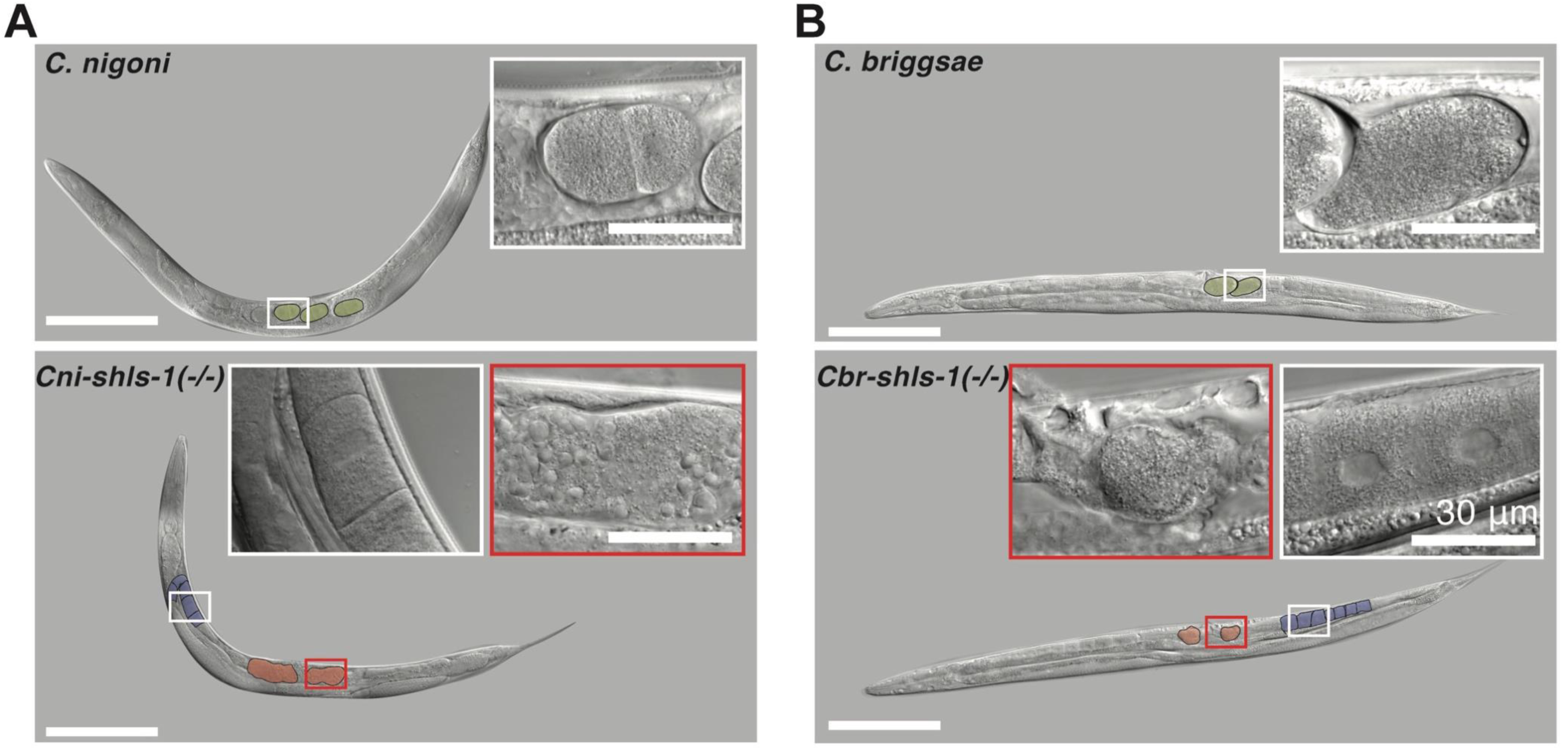
Malformation of embryos in *shls-1* homozygous mutant females. **(A-B)** DIC images of WT (top) and *shls-1(-/-)* mutant (bottom) adult females of *C. nigoni* (A) and *C. briggsae* (B). WT embryos are pseudo-colored yellow, deformed *shls-1* mutant embryos pink, and intact mutant oocytes purple. Insets show magnified views of intact embryos and oocytes (white rectangles), and deformed embryos (red rectangles). Scale bar: 200 μm (original); 30 μm (insets).

**Figure S5.**
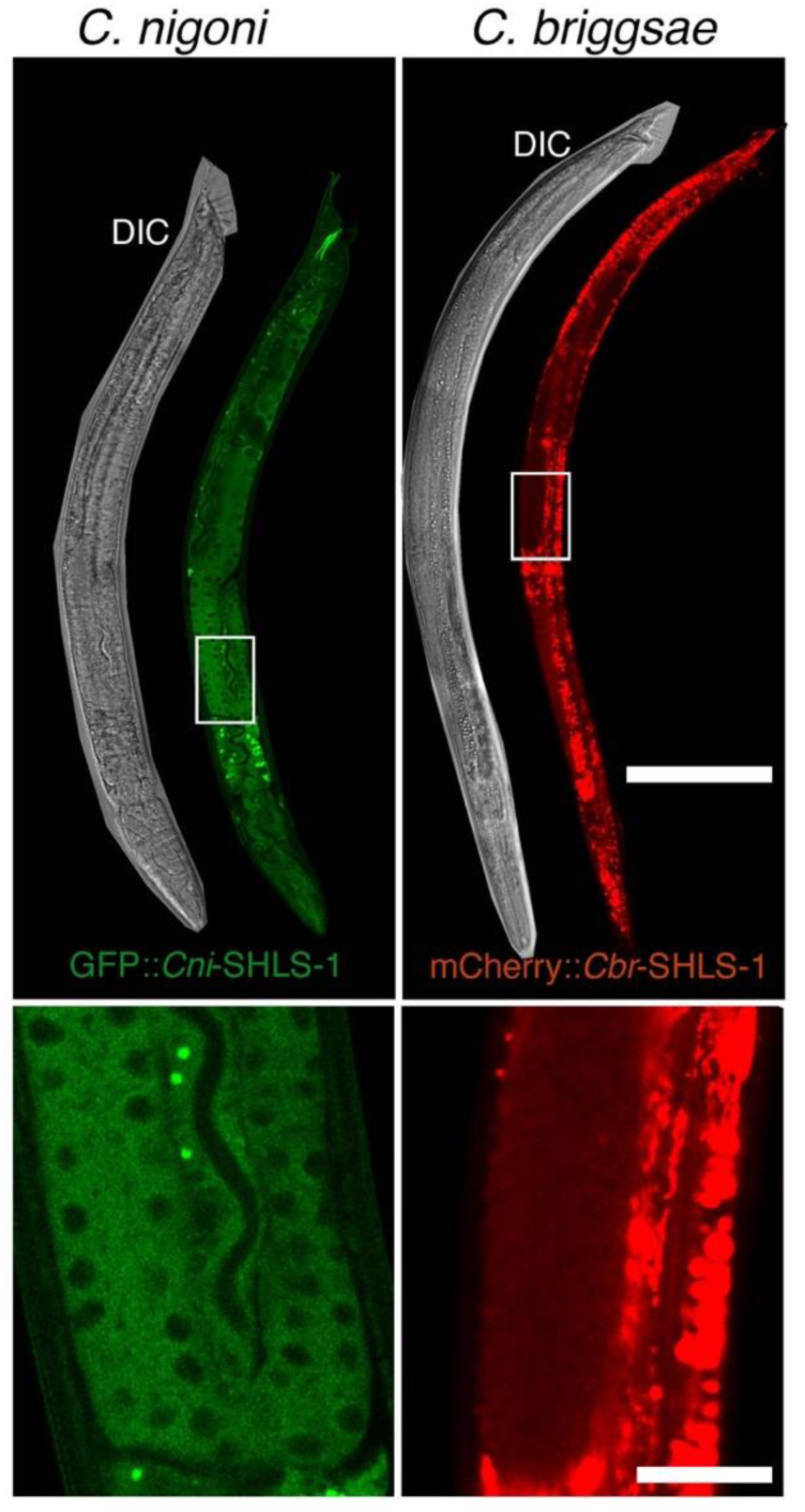
Expression of SHLS-1 in adult male of *C. nigoni* (left) and *C. briggsae* (right) The *C. nigoni* and *C. briggsae* male carry N-terminally tagged GFP::*Cni-*SHLS-1 and mCherry::*Cbr-*SHLS-1, respectively as shown in Fig. 2D. Magnified images highlight expression in male germlines. Scale bar: 100µm (original); 20µm (magnified).

**Figure S6.**
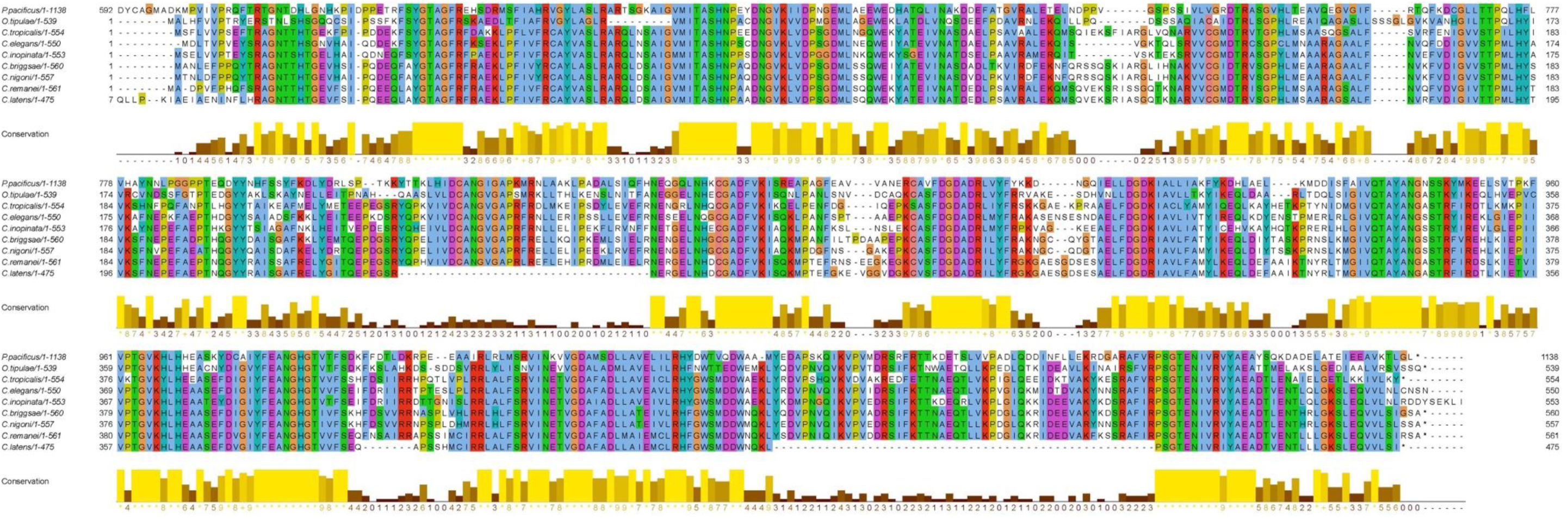
Conservation of SHLS-1 proteins across nematodes. Shown is the multiple sequence alignment of SHLS-1 proteins from seven *Caenorhabditis* nematodes and two distantly related nematodes, i.e., *Pristionchus Pacificus* and *Oscheius Tipulae*. Note that greater divergence from *P. pacificus* and *C. latens* may be due to inaccurate gene annotation. Conservation scores are shown at the bottom. Color code follows the ClustalX style.

**Figure S7.**
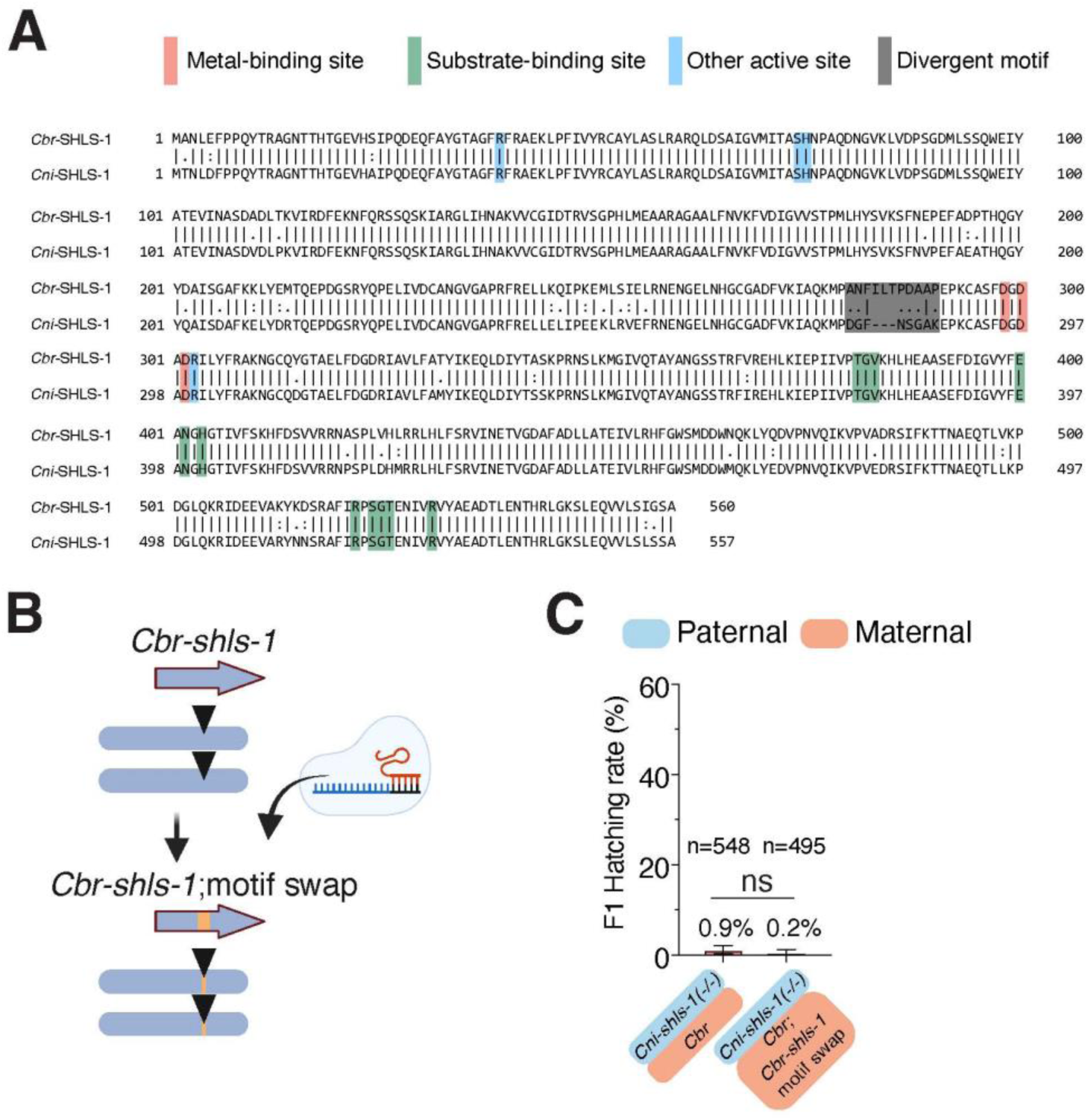
The divergent motif between *Cni-*SHLS-1 and *Cbr-*SHLS-1 is not responsible for hybrid lethality. **(A)** Pairwise protein sequence alignment of *Cbr-*SHLS-1 and *Cni-*SHLS-1. Active sites including metal-binding (pink), substrate-binding (green) and others (blue), are highlighted (see Fig. 3A). The highly divergent motif is shaded with black. **(B)** Schematic showing the replacement of the divergent motif in *C. briggsae* with the sequence from *C. nigoni* using CRISPR/Cas9. **(C)** Comparison of hatching rates for F1 hybrids from *Cni-shls-1(-/-)* mutant fathers and *C. briggsae* mothers or *C. briggsae* mothers with the replaced motif. *P*=0.2216, Fisher’s exact test; error bars: 95% confidence interval.

**Figure S8.**
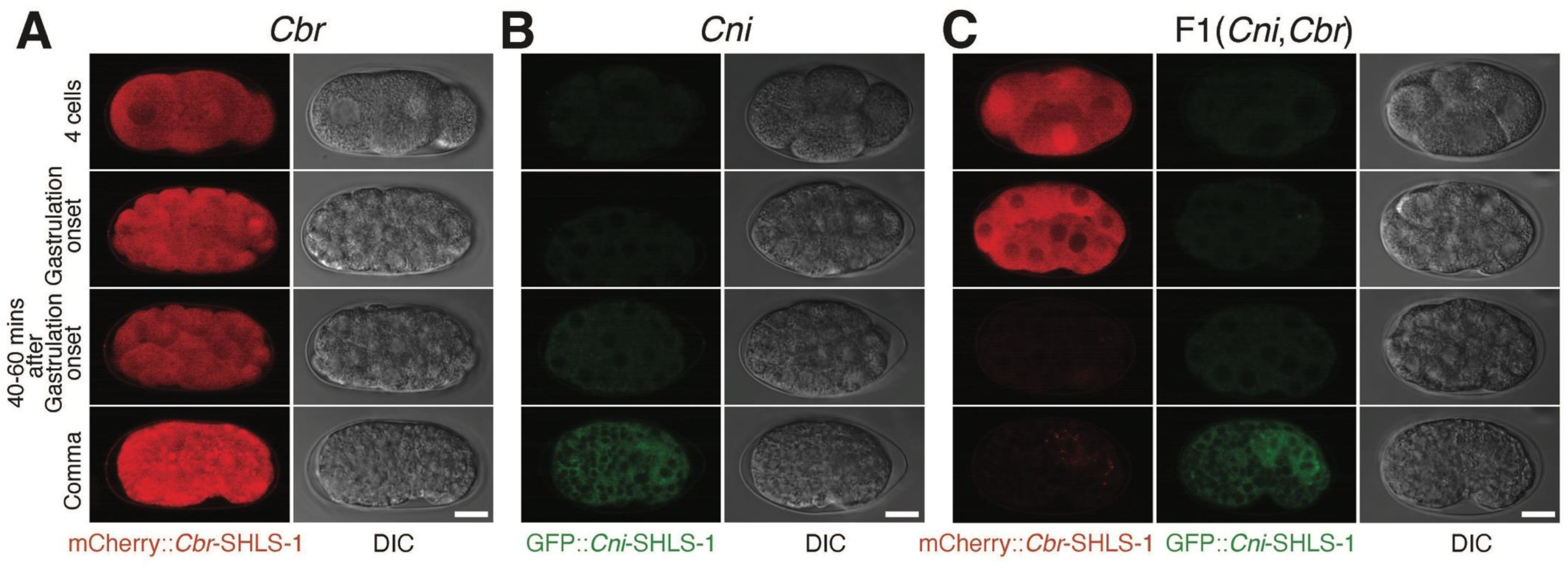
Rapid degradation of *C. briggsae* SHLS-1 in F1 hybrids. Representative fluorescent images of *Cbr-*SHLS-1 or *Cni-*SHLS-1 expression (left) and corresponding DIC images (right) in embryos of *C. briggsae* carrying heterozygous *mCherry::Cbr-shls-*1 (A), *C. nigoni* carrying heterozygous *gfp::Cni-shls-1* (B) or F1 hybrids from *C. nigoni* fathers carrying homozygous *gfp::Cni-shls-1* and *C. briggsae* mothers carrying homozygous *mCherry::Cbr-shls-1* (C) across four developmental stages: 4-cell, gastrulation onset, 40-60 mins after gastrulation onset and comma stage. Images are from Fig. 3C and D, supplemented with DIC images. Scale bar, 10µm.

**Figure S9.**
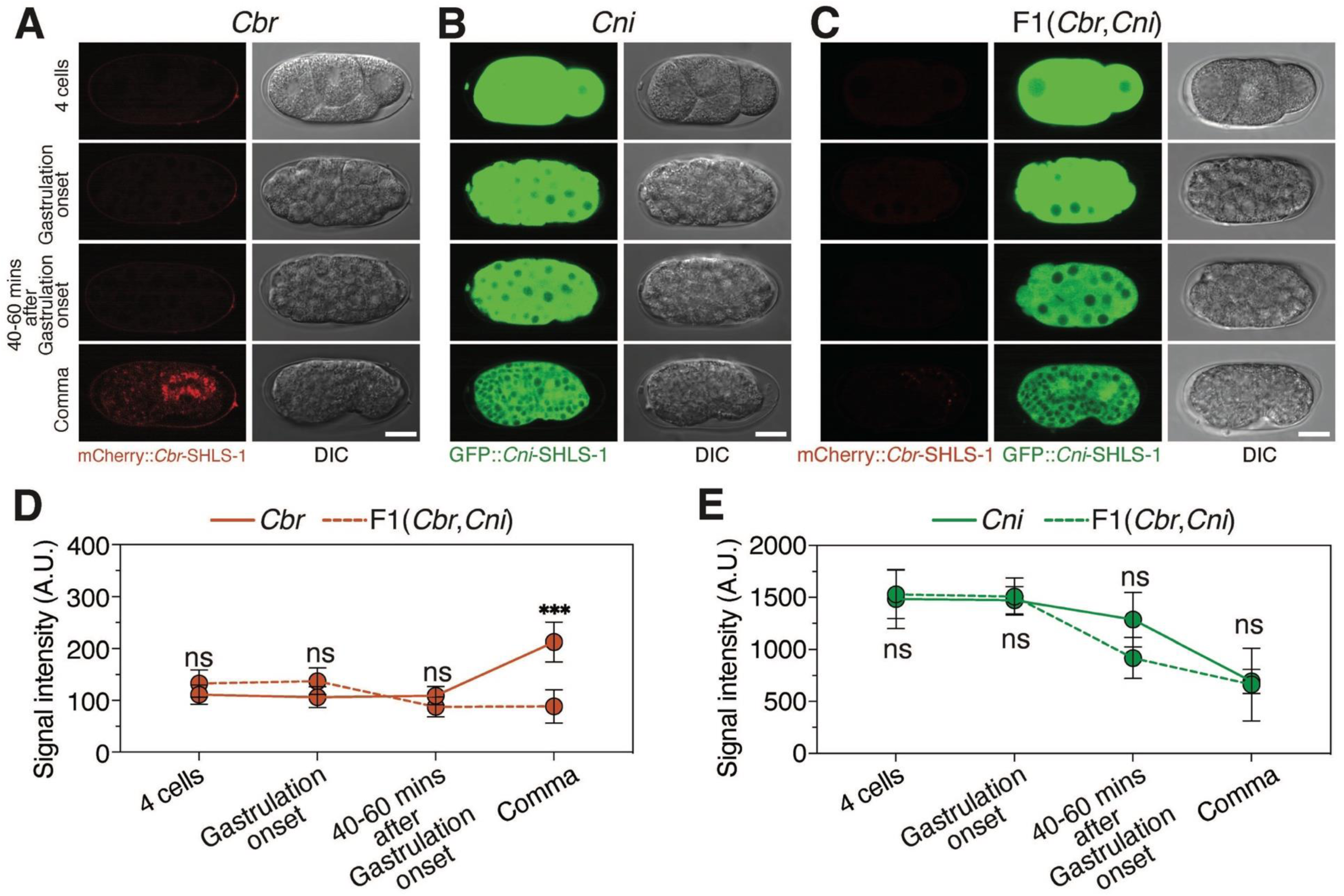
Specific degradation of *Cbr-*SHLS-1 in F1 hybrids from *C. briggsae* fathers and *C. nigoni* mothers. **(A-C)** Representative images showing *Cbr-*SHLS-1 expression in *C. briggsae* embryos carrying heterozygous *mCherry::Cbr-shls-1* (A), *Cni-* SHLS-1 expression *C. nigoni* embryos carrying heterozygous *gfp::Cni-shls-1* (B) and both expressions in the F1 hybrid embryos from *C. nigoni* mothers carrying homozygous *gfp::Cni-shls-1* and *C. briggsae* fathers carrying homozygous *mCherry::Cbr-shls-1* (C) across the same developmental stages as in Fig 3C and D. Genotype details: *Cbr* embryos are *mCherry::Cbr-shls-1(+/-)* from *mCherry::Cbr-shls-1(+/+)* fathers and WT mothers; *Cni* embryos are *gfp::Cni-shls-1(+/-)* embryos from WT fathers and *gfp::Cni-shls-1(+/+)* mothers; F1(*Cbr,Cni*) embryos are *mCherry::Cbr-shls-1(+)/gfp::Cni-shls-1(+)* from *Cbr-shls-1(+/+)* fathers and *gfp::Cni-shls-1(+/+)* mothers. Scale bar, 10µm. **(D-E)** Quantification of the mCherry (D) or GFP intensity (E) in hybrid F1 embryos from *C. nigoni* mothers and *C. briggsae* fathers compared to *C. briggsae* (mCherry) or *C. nigoni* (GFP) embryos at each stage shown in (A-C). n=5 for each stage; \*\*\**P=*0.00054, unpaired t-test; Error bar: s.e.m; A.U., Arbitrary units.

**Figure S10.**
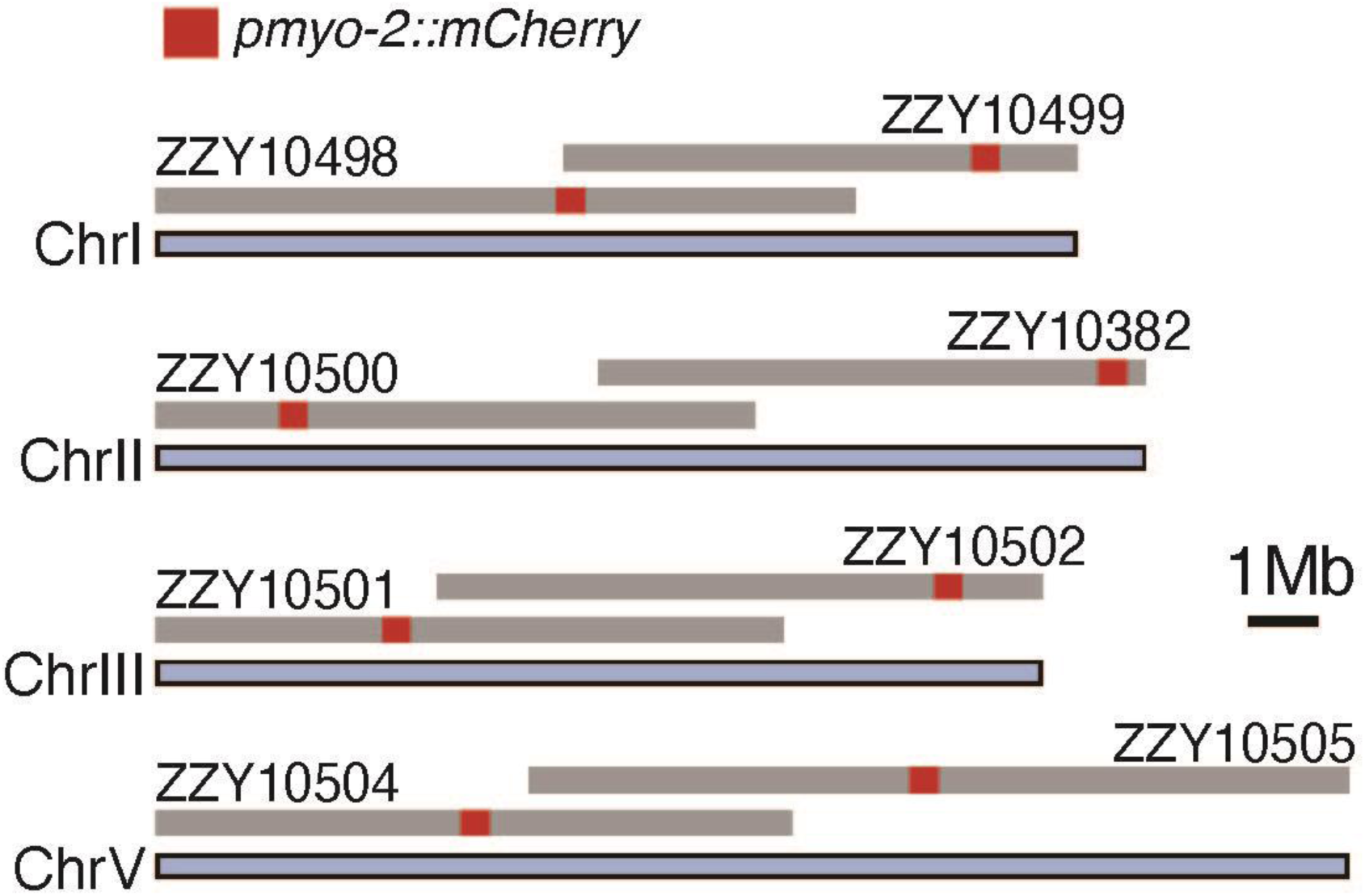
Sizes of introgressions on autosomes other than Chr IV in IR strains with dual introgression fragments (see Fig. 4D) Introgression fragments are drawn in scale as gray horizontal bars above *C. briggsae* chromosomes. Also shown are the genomic position of each integrated mCherry marker, i.e., *pmyo-2::mCherry*.

**Figure S11.**
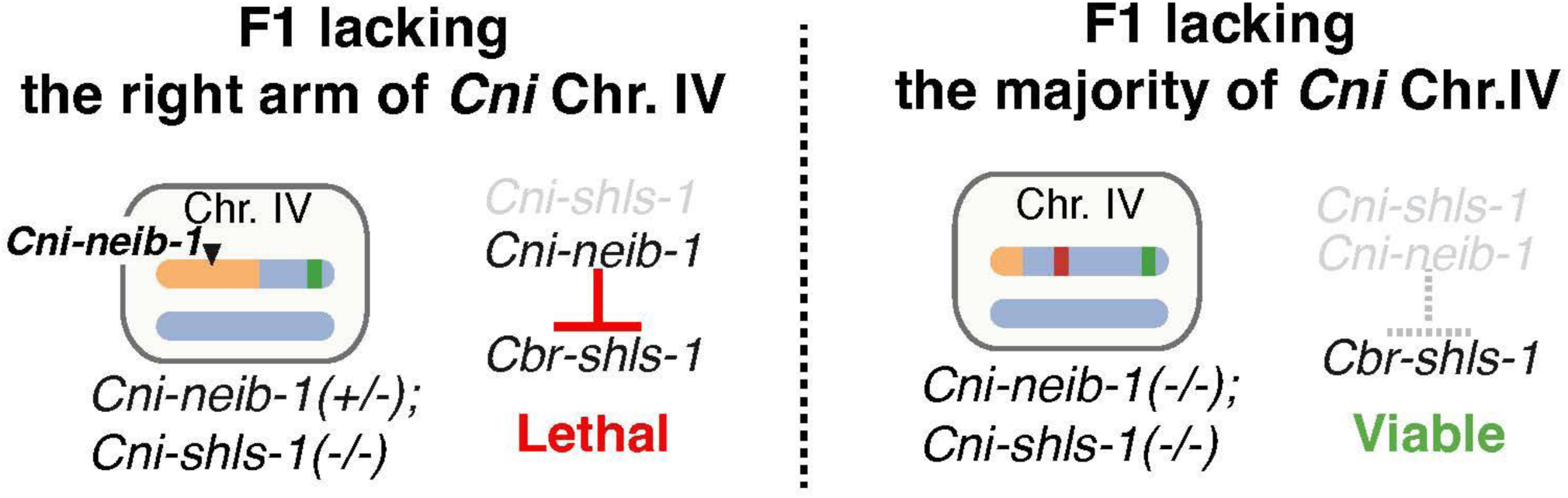
Schematic illustrating rationale for rescuing hybrid lethality in F1 hybrids lacking most of *C. nigoni* Chr. IV. *Cni-neib-1* negatively interact with *Cbr-shls-1* in F1 hybrids lacking *C. nigoni* Chr. IV right arm including *Cni-shls-1* (left). If *Cni-neib-1* is located on the left half of *C. nigoni* Chr. IV, an extended *C. briggsae* introgression removing most of *C. nigoni* Chr. IV, including *Cni-neib-1* in the F1 hybrids, is expected to rescue the lethality although *Cni-shls-1* is also missing (right).

**Figure S12.**
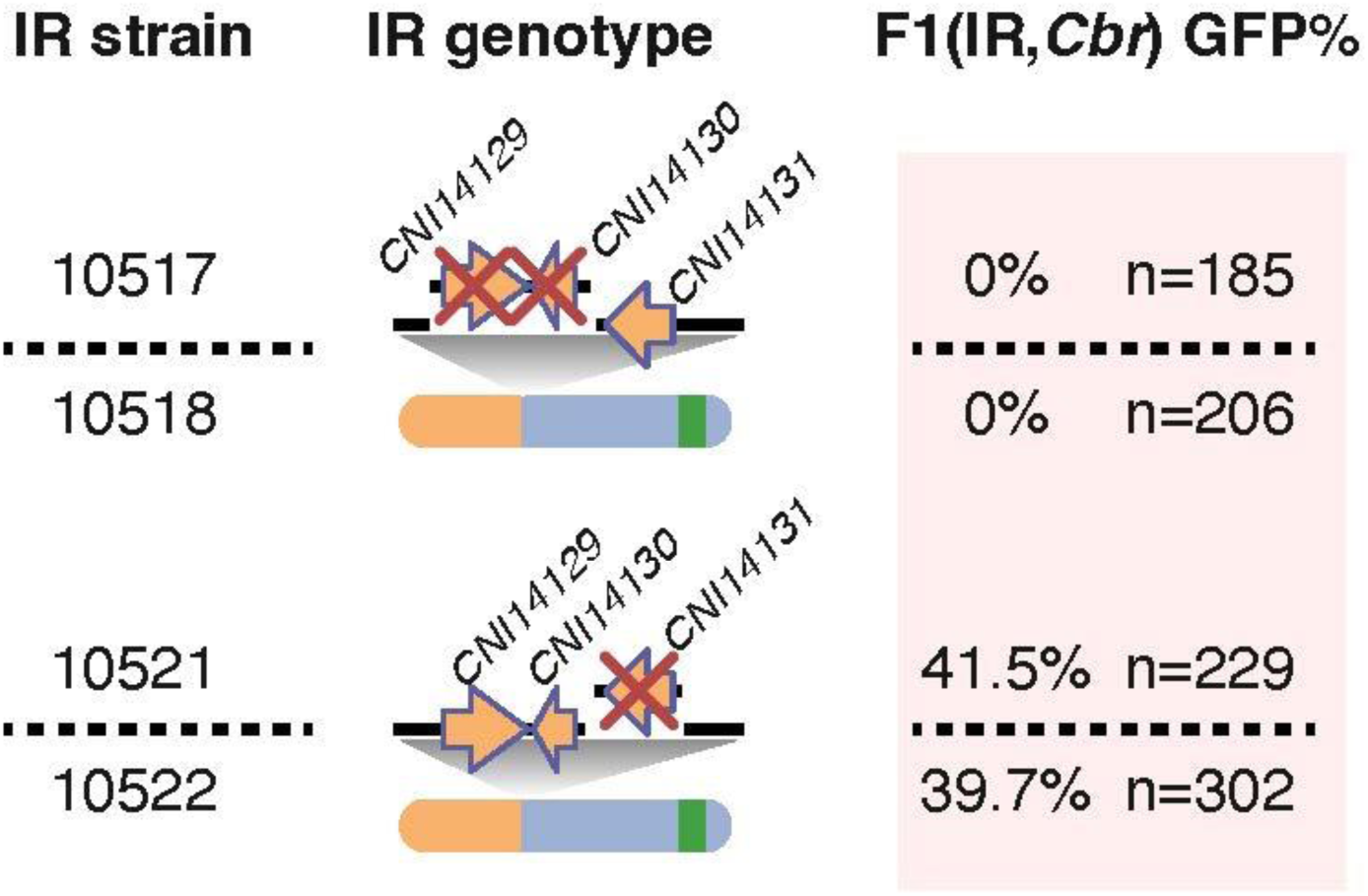
Cloning of *Cni-neib-1* through deletion(s) of candidate gene(s) in the ∼37kb mapped interval (see Fig. 4E) Shown are the percentages of GFP+ hybrid F1 adults from *C. briggsae* mothers and IR strain fathers with genomic deletion(s) of both *CNI14129* and *CNI14130* (top), or *CNI14131* only (bottom) on *C. nigoni* Chr. IV carrying the *C. briggsae* introgression. Deletions were generated in IR strain ZZY10481 (see Fig. 4E). Genotype schematics for *Cni-*Chr. IVs with *C. briggsae* introgressions are shown. Two independent lines were generated for each type of deletion.

**Figure S13.**
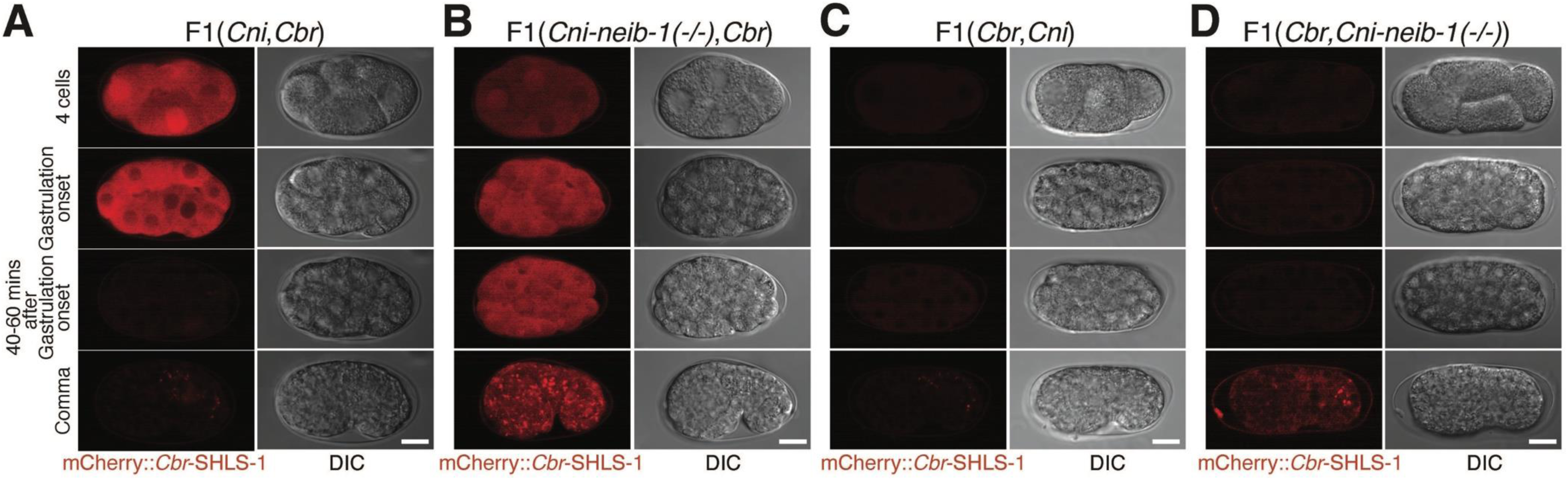
Deletion of *Cni-neib-1* prevents degradation of *Cbr-*SHLS-1 in F1 hybrids. **(A-D)** Representative fluorescent images of *Cbr-*SHLS-1 expression (left) and corresponding DIC images (right) in hybrid F1 embryos from *C. briggsae* mothers carrying homozygous *mCherry::Cbr-shls-1* with *C. nigoni* fathers (A) or *Cni-neib-1(-/-)* mutant fathers (B), and from the same transgenic *C. briggsae* fathers with *C. nigoni* mothers (C) or *Cni-neib-1(-/-)* mutant mothers (D), across the same developmental stages shown in Fig 4G and I. Images are from Fig. 4G and I, supplemented with DIC images. Scale bar, 10µm

**Figure S14.**
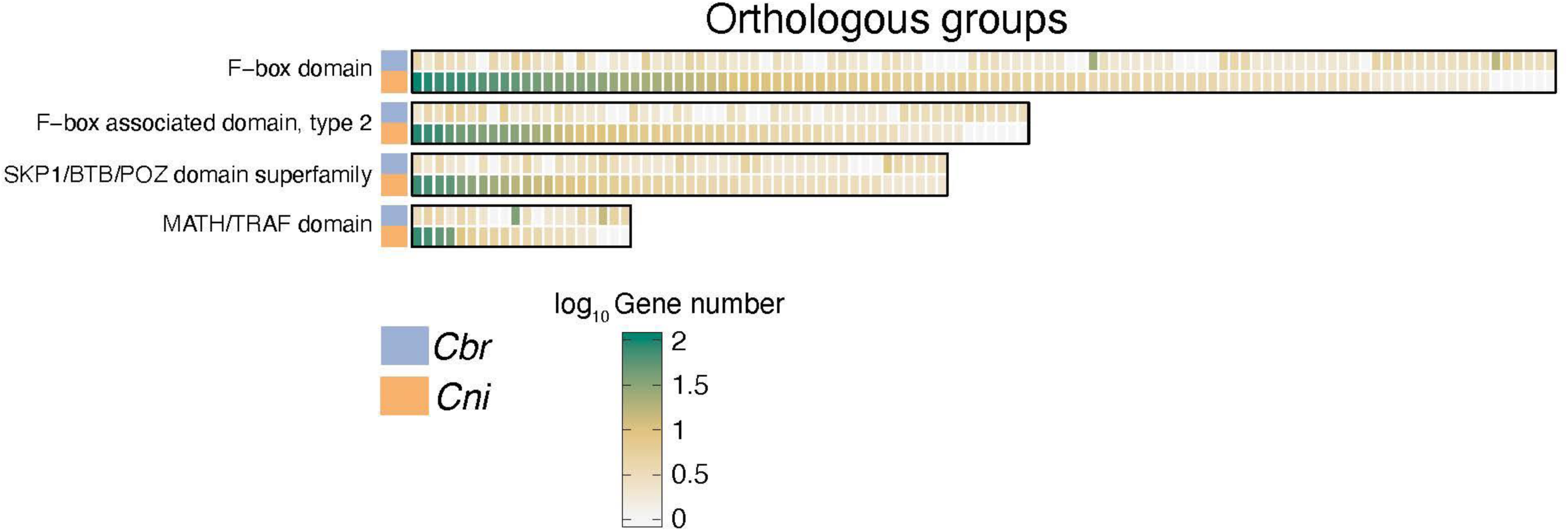
Comparison of gene numbers between *C. nigoni* and *C. briggsae* in the orthologous groups (shown as vertical bars) belonging to *C. nigoni*-expanded protein domains (see Fig. 5B) Protein domains are shown on the left. Gene number is shown in log10 scale as indicated. Note that extensive expansion of most orthologous groups in *C. nigoni* compared to *C. briggsae*.

**Figure S15.**
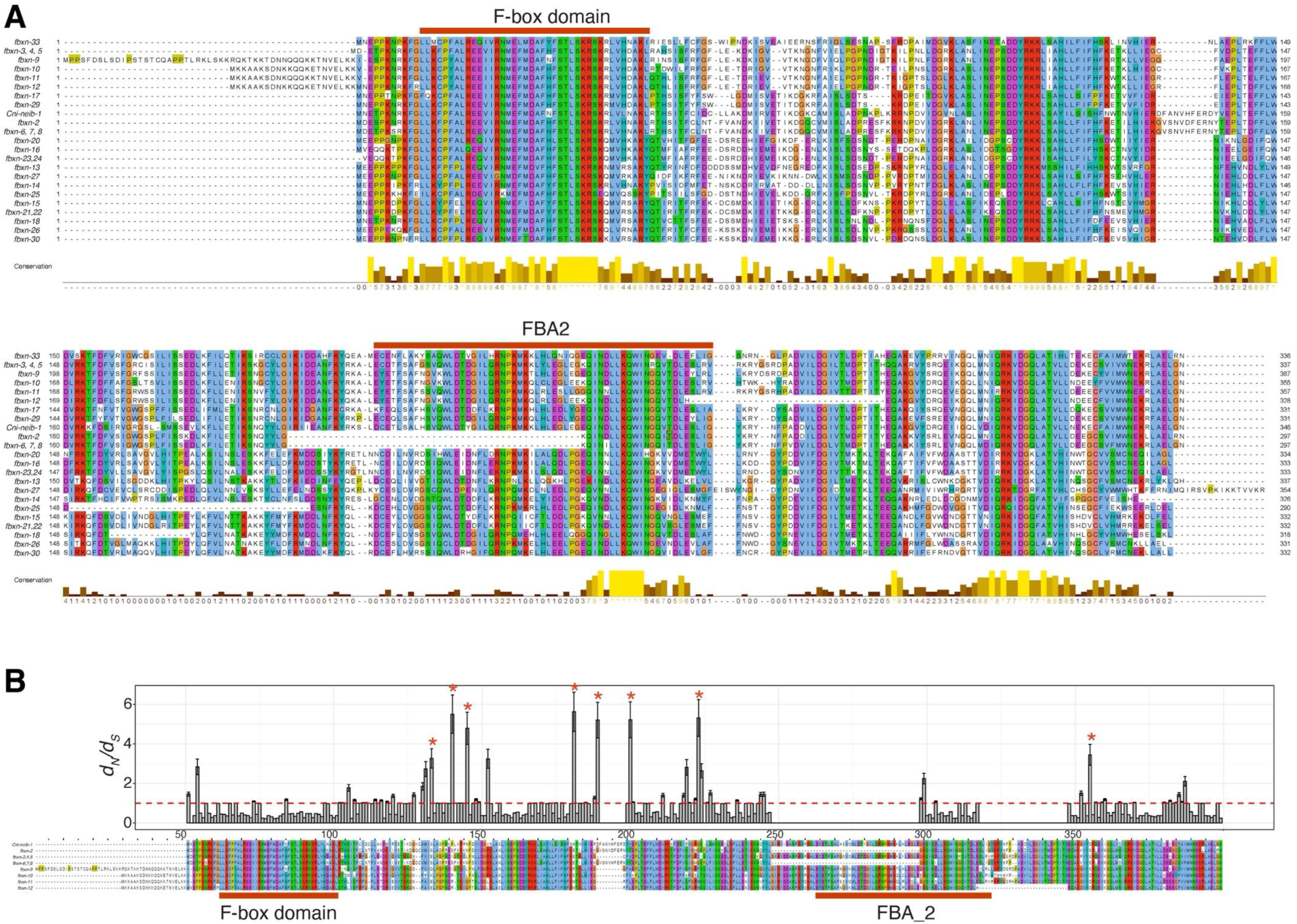
*fbxn* genes are rapidly evolving and highly dynamic. **(A)** Protein sequence alignment of paralogues in the *fbxn* gene family. *fbxn-19, −28, −31,* and *-32* are excluded because they underwent more extensive gene structural changes than the others and might be pheudogenized. Conservation scores are shown at the bottom. The color code follows Clustal X style. The F-box domain and F-box associated domain 2 (FBA2) are indicated by red bars. **(B)** Maximum-likelihood *d_N_/d_S_* values (top) and the protein sequence alignment (bottom) for *Cni-neib-1* and its most related *fbxn* paralogues (*fbxn-2* to *fbxn-12*). The F-box domain and FBA2 domain are highlighted by red bars. Bar plots above the alignment denote the values of estimated *d_N_/d_S_* for each gap-free alignment column. Sites with a high probability of positive selection (*P*>0.8) are labeled with red asterisks. Red dashed lines indicate a *d_N_/d_S_* value of 1.

**Figure S16.**
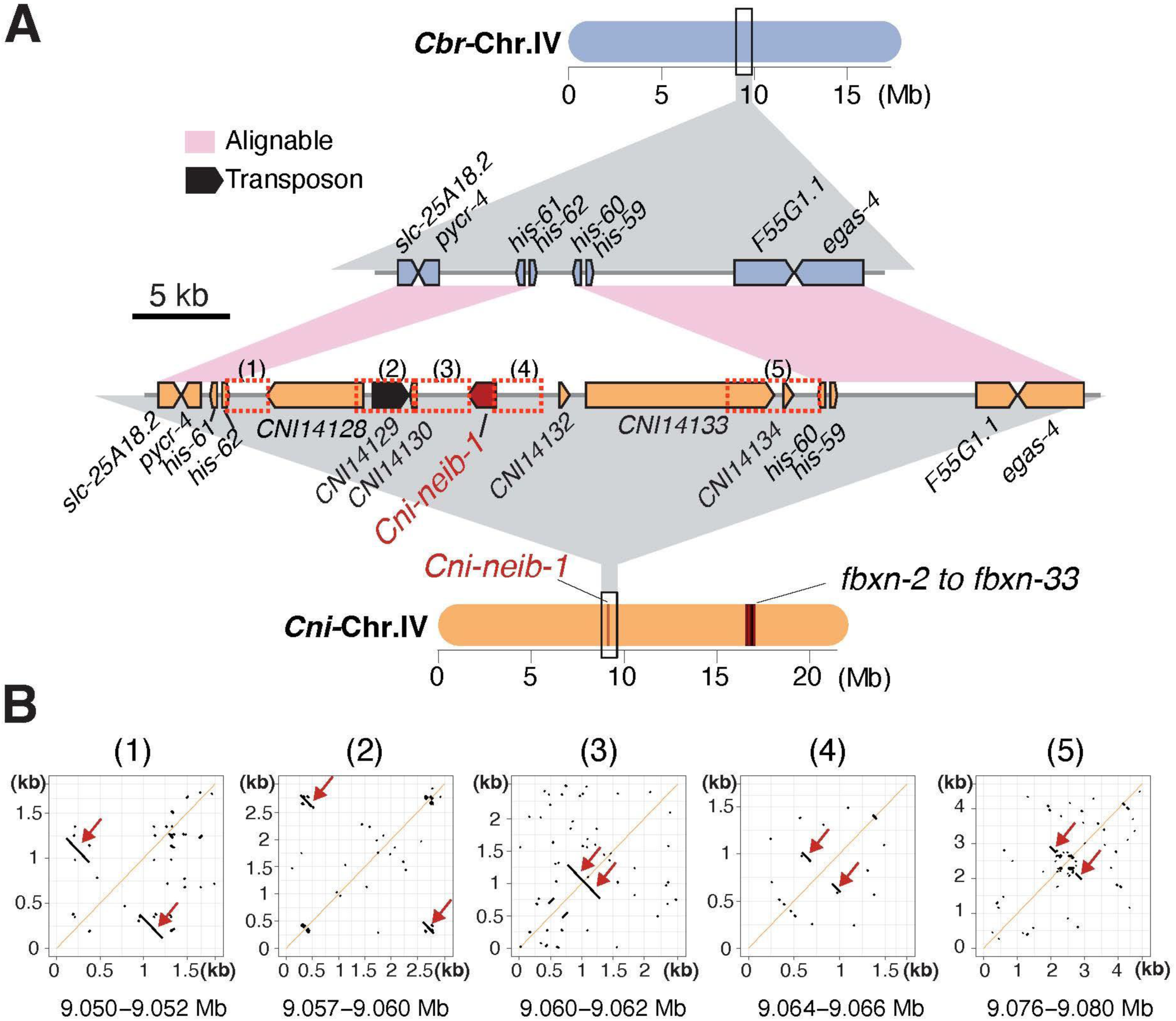
*Cni-neib-1* resides in a *C. nigoni*-specific genomic region characterized by frequent DNA inversions, duplications and transpositions. **(A)** Comparison of syntenic regions between *C. briggsae* and *C. nigoni* containing *Cni-neib-1,* which was translocated to the center of Chr. IV. Red dashed boxes highlight five regions enriched with inverted repeats, direct repeats and short duplicates. Note that an intact transposon is located adjacent to the 5’ end of *Cni-neib-1*. *C. elegans* gene names are used for all orthologous genes for simplicity. **(B)** Dot plots of genomic sequences from each highlighted region shown in (A). Red arrowheads mark the longest inverted repeats within each region.

**Figure S17.**
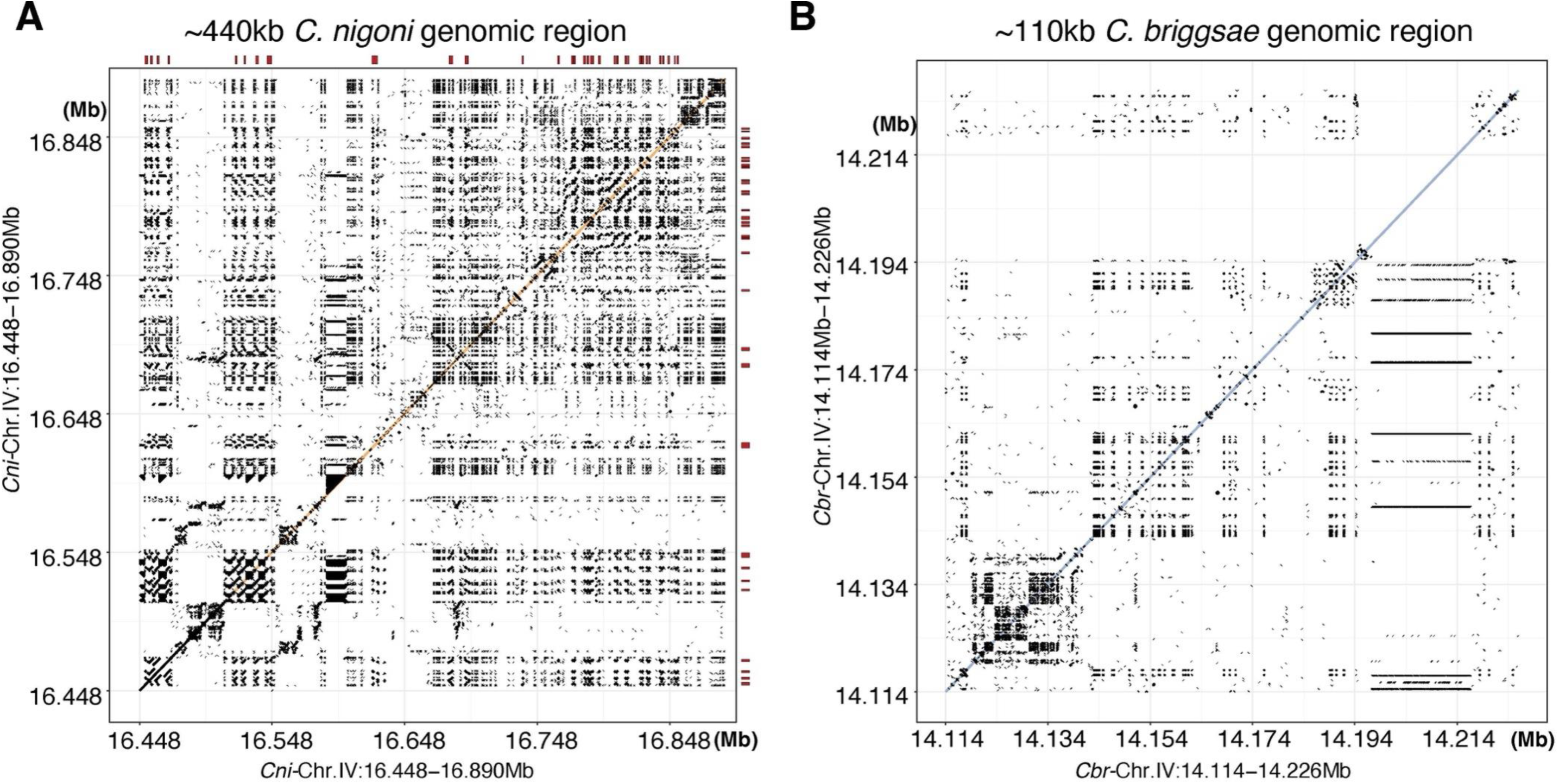
The ∼440kb *C. nigoni* genomic region (see Fig. 5E) is enriched with tandem repeats flaking *fbxn* genes. **(A)** Dot plot of the ∼440kb *C. nigoni* genomic region (see Fig. 5E) containing the 32 *fbxn* genes and *Cni-shls-1*. Abundant tandem simple repeats are distributed across the entire region and located in proximity to the *fbxn* genes. Positions of the *fbxn* genes are indicated in scale as red bars. **(B)** Dot plot of the syntenic ∼110kb *C. briggsae* genomic region containing *Cbr-shls-1* (see Fig. 5E). Note the significantly reduced density of repeats in the *C. briggsae* syntenic region.

**Figure S18.**
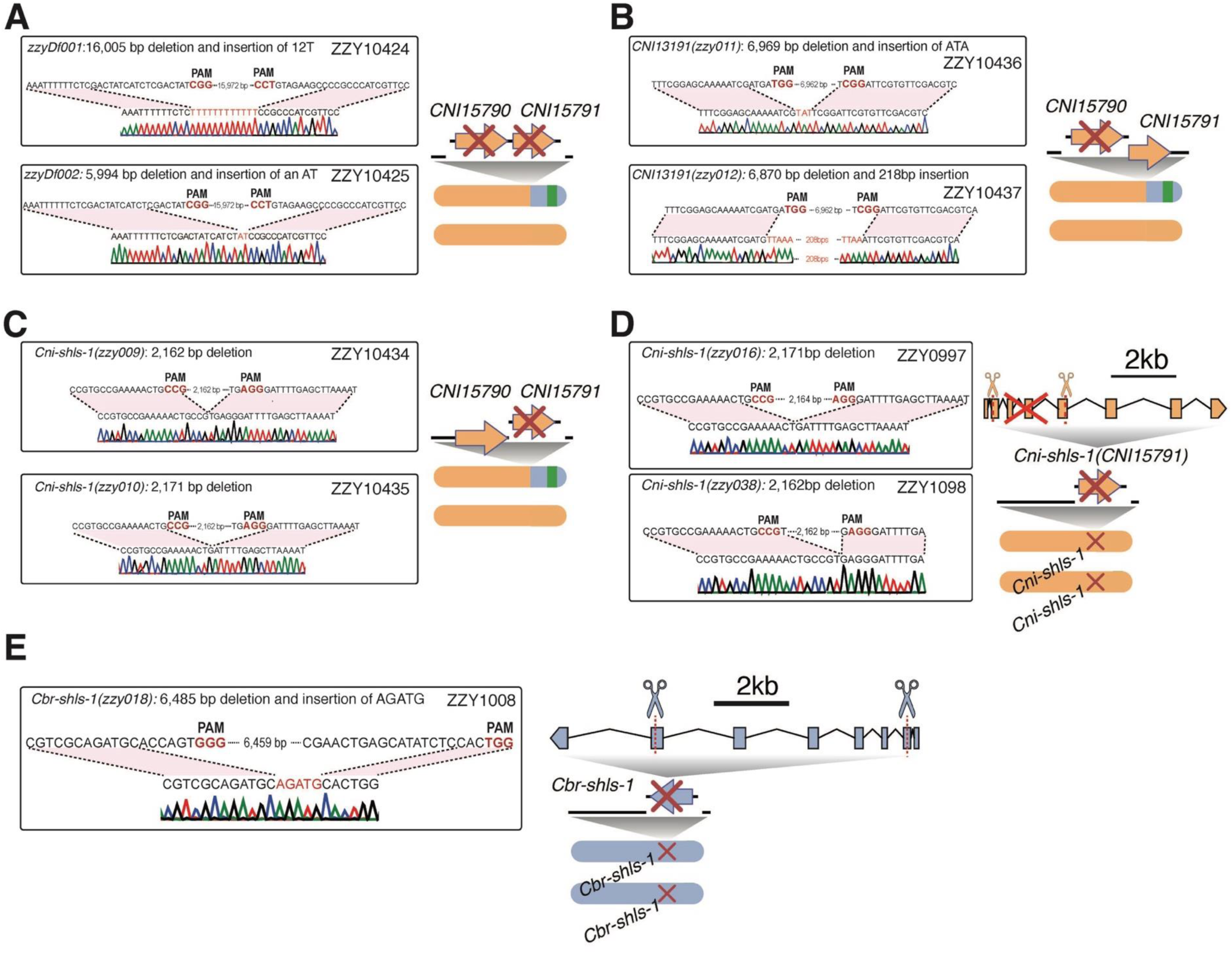
Confirmation of deletion alleles generated for the identification of *Cni-shls-1*. **(A-E)** Chromatograms (left) and schematics (right) of deletion alleles (indicated with “X”) generated using dual gRNAs targeting the *C. nigoni* chromosomes with *C. briggsae* introgression fragments in IR strains (A-C, see Fig. S2), or targeting chromosomes of WT *C. nigoni* (D, see Fig. 2B and Fig. S4A) or *C. briggsae* (E, see Fig. S4B). Strain names, deletion allele descriptions, PAM sequences, and sequences of deletion junctions confirmed by Sanger sequencing are provided. Two independent alleles were generated for each deletion.

**Figure S19.**
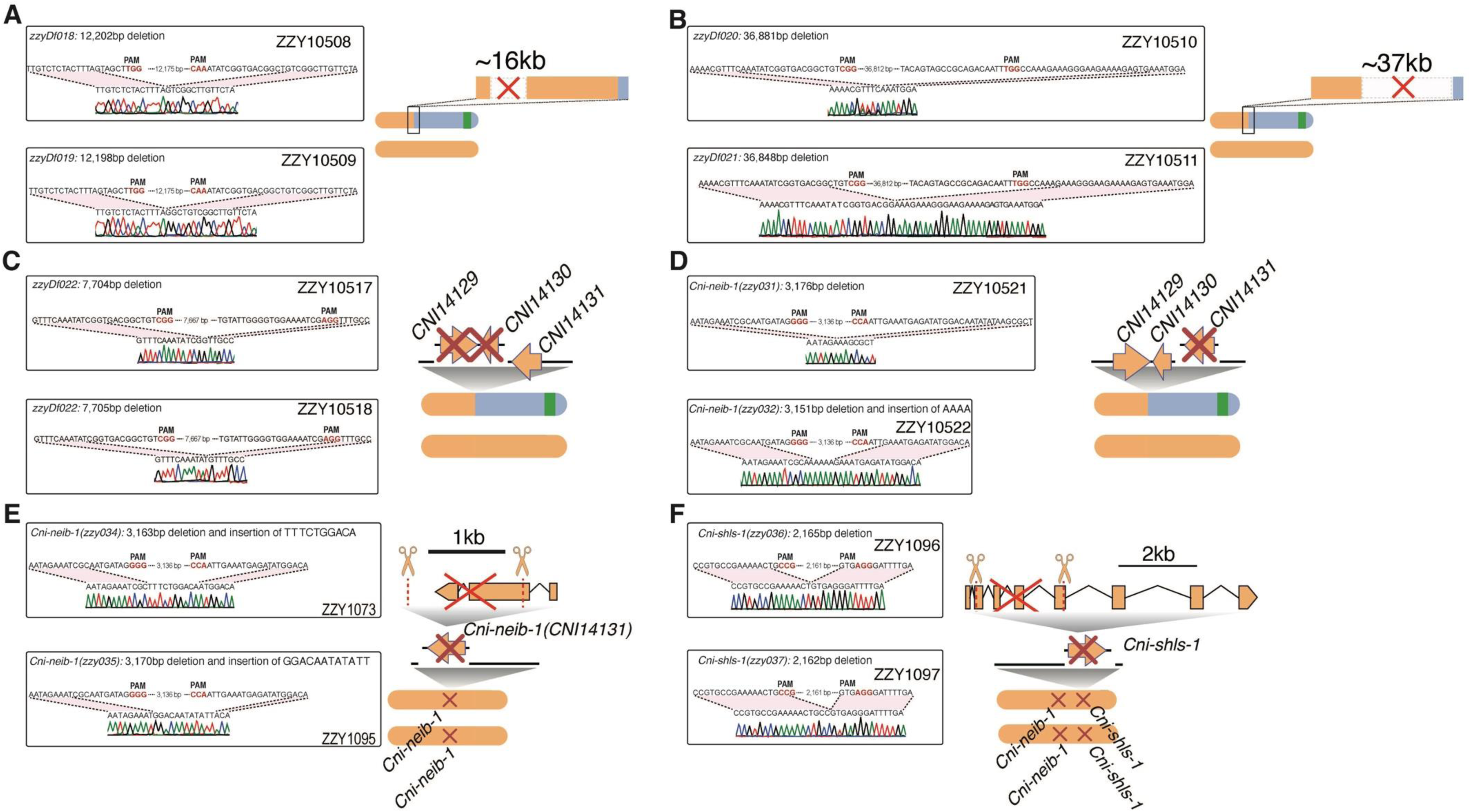
Confirmation of deletion alleles generated for the identification of *Cni-neib-1*. **(A-F)** Chromatograms (left) and schematics (right) of the deletion alleles (indicated with “X”) generated using dual gRNAs targeting the *C. nigoni* chromosomes with *C. briggsae* introgression fragments in IR strains (A-D, see Fig. 4E and Fig. S12), or targeting chromosomes of WT *C. nigoni* (E and F, see Fig. 4F). Strain names, deletion allele descriptions, PAM sequences and deletion junctions confirmed by Sanger sequencing are provided. *Cni-shls-1* and *Cni-neib-1* double mutant (ZZY1096 and ZZY1097) were generated by knocking out *Cni-shls-1* in the *Cni-neib-1* homozygous mutant, i.e., ZZY1073 shown in (E). Two independent alleles were generated for each deletion.

**Table S2.** List of *fbxn* genes.

| Gene ID | F-box Gene name | Pseudogene | Remarks |
| --- | --- | --- | --- |
| *CNI14131* | *Cni-neib-1* | No |  |
| *CNI15784* | *fbxn-2* | No |  |
| *CNI15785* | *fbxn-3* | No |  |
| *CNI15786* | *fbxn-4* | No |  |
| *CNI15787* | *fbxn-5* | No | *fbxn-3,4,5* are tandem duplicates |
| *CNI15793* | *fbxn-6* | No |  |
| *CNI15794* | *fbxn-7* | No |  |
| *CNI15795* | *fbxn-8* | No | *fbxn-6,7,8* are tandem duplicates |
| *CNI15796* | *fbxn-9* | No |  |
| *CNI15805* | *fbxn-10* | No |  |
| *CNI15812* | *fbxn-11* | No |  |
| *CNI15814* | *fbxn-12* | No |  |
| *CNI15822* | *fbxn-13* | No |  |
| *CNI15828* | *fbxn-14* | No |  |
| *CNI15832* | *fbxn-15* | No |  |
| *CNI15833* | *fbxn-16* | No |  |
| *CNI15836* | *fbxn-17* | No |  |
| *CNI15837* | *fbxn-18* | No |  |
| *CNI15838* | *fbxn-19* | Yes | local duplication of exon 1 and 2 |
| *CNI15840* | *fbxn-20* | No |  |
| *CNI15842* | *fbxn-21* | No | *fbxn-21,23* are tandem duplicates |
| *CNI15843* | *fbxn-22* | No | *fbxn-22,24* are tandem duplicates |
| *CNI15844* | *fbxn-23* | No | *fbxn-21,23* are tandem duplicates |
| *CNI15845* | *fbxn-24* | No | *fbxn-22,24* are tandem duplicates |
| *CNI15847* | *fbxn-25* | No |  |
| *CNI15848* | *fbxn-26* | No |  |
| *CNI15850* | *fbxn-27* | No |  |
| *CNI15851* | *fbxn-28* | Yes | transposon insertion |
| *CNI15854* | *fbxn-29* | No |  |
| *CNI15855* | *fbxn-30* | No |  |
| *CNI15856* | *fbxn-31* | Yes | transposon insertion |
| *CNI15857* | *fbxn-32* | Yes |  |
| *CNI15858* | *fbxn-33* | No |  |

**Table S4.**
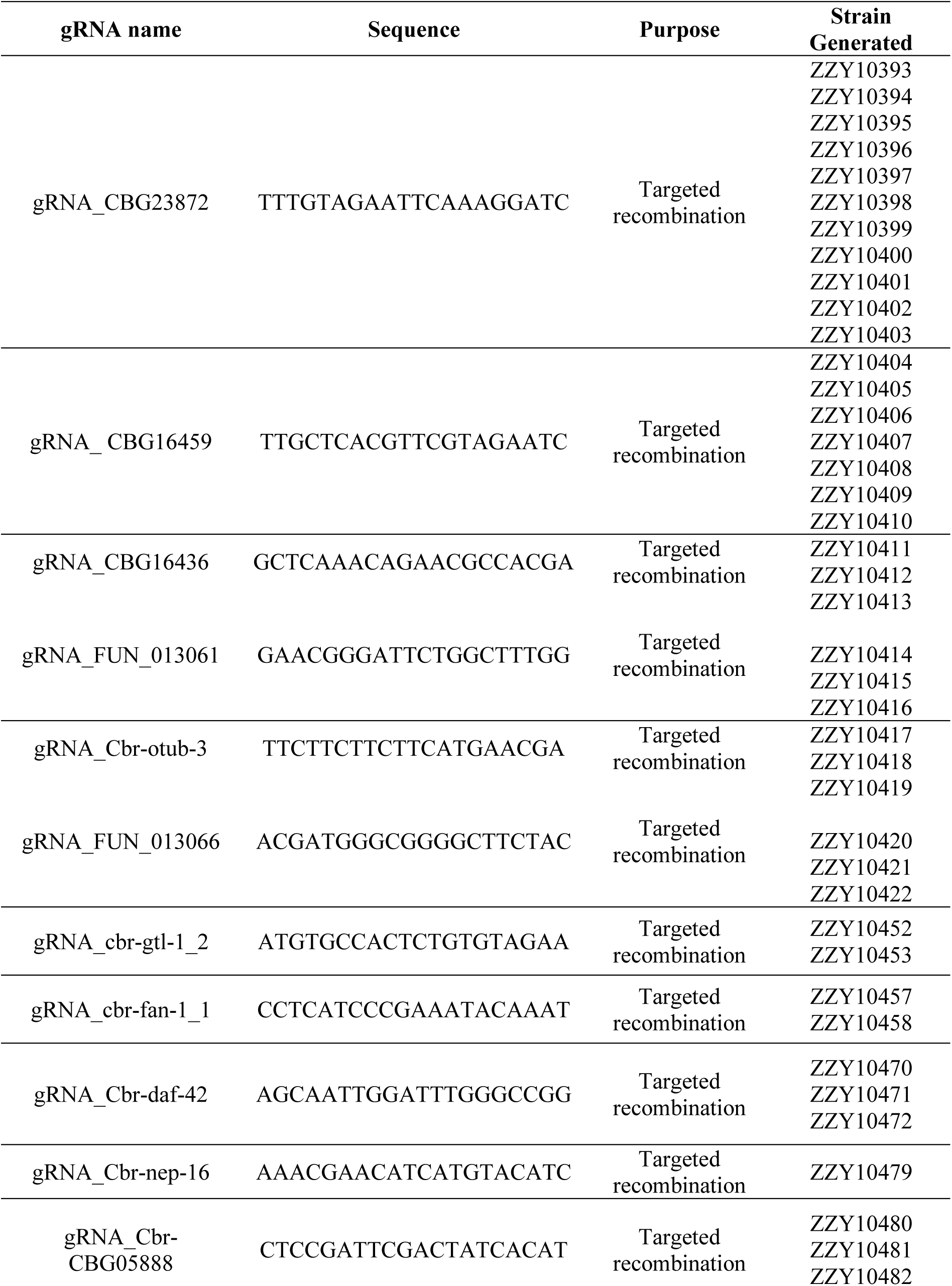
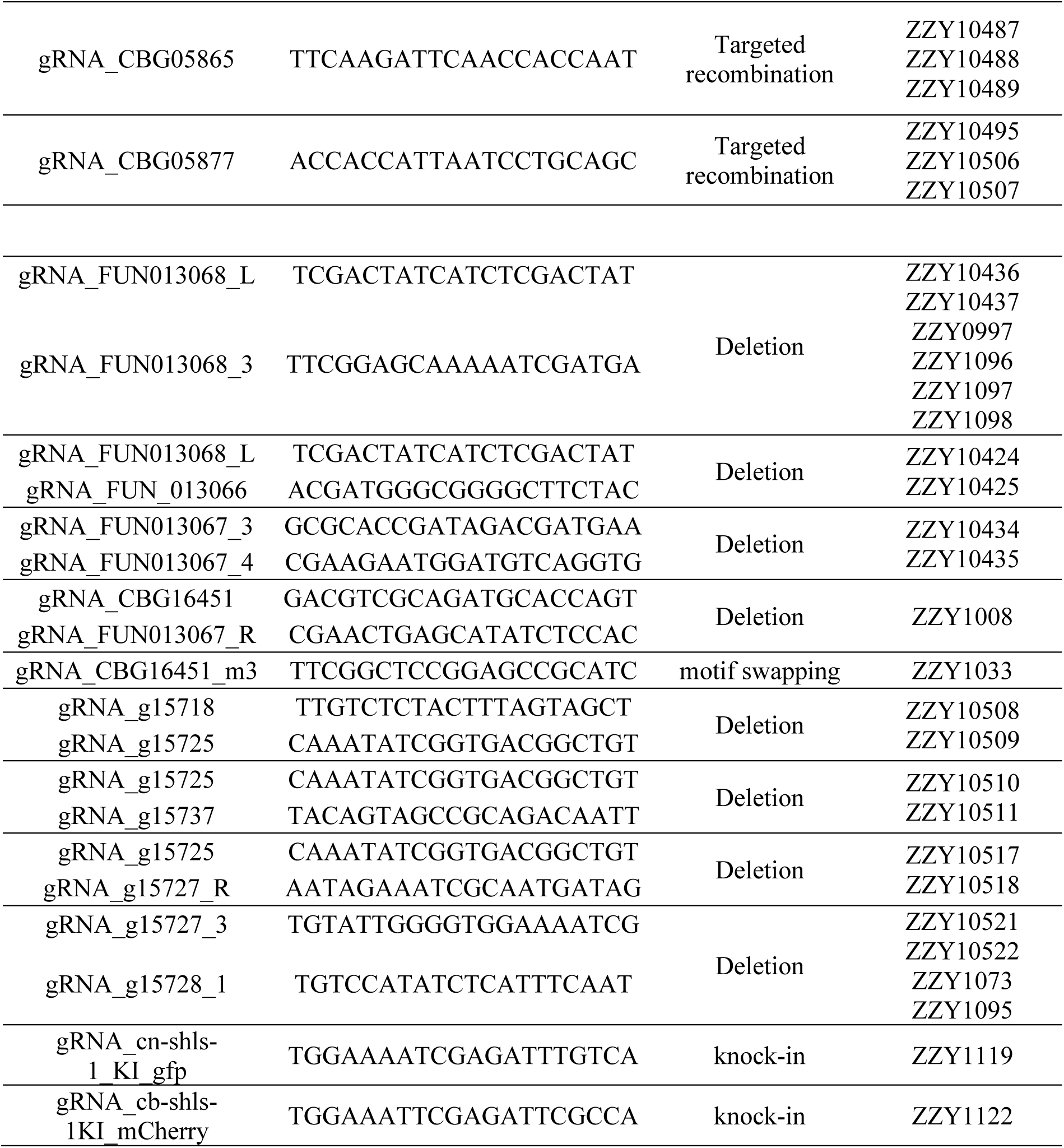
List of gRNAs sequences for targeted recombination and genome editing.

**Table S5.**
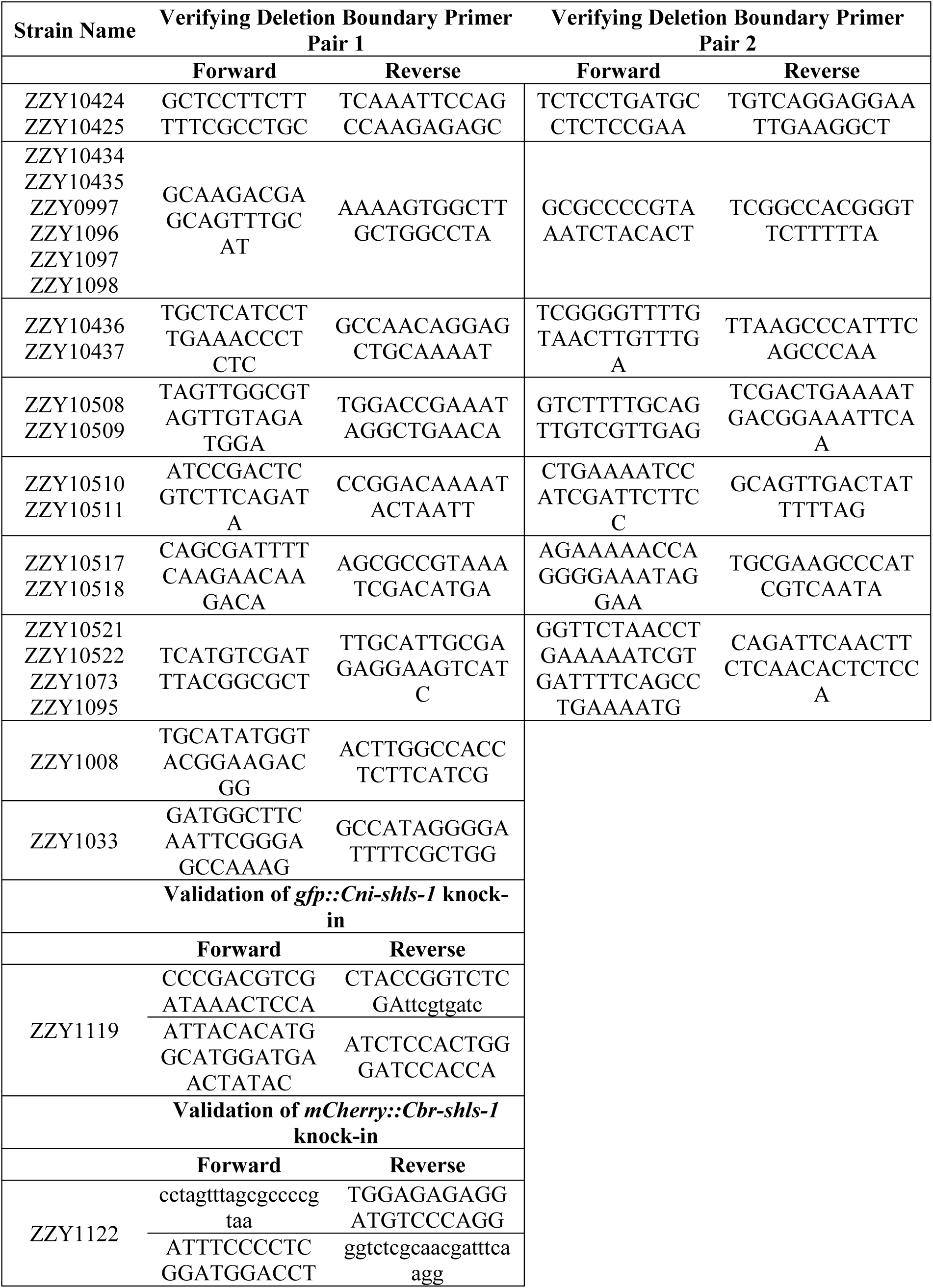
List of primers used for amplifying the deletion boundaries.

**Table S6.** List of genome-edited and transgenic strains generated in this study.

| Strain Name | Genotype | Allele Name |
| --- | --- | --- |
| **Deletion and Knockout Strain** |  |  |
| ZZY10424 | *C. nigoni*, zzyDf001 IV, zzyIR10417 (IV, AF16>JU1421), zzyIs0734[Cbr-myo-2p::GFP; NeoR] IV | zzyDf001 |
| ZZY10425 | *C. nigoni*, zzyDf002 IV, zzyIR10417 (IV, AF16>JU1421), zzyIs0734[Cbr-myo-2p::GFP; NeoR] IV | zzyDf002 |
| ZZY10434 | *C. nigoni*, Cni-shls-1(zzy009) IV, zzyIR10417 (IV, AF16>JU1421), zzyIs0734[Cbr-myo-2p::GFP; NeoR] IV | Cni-shls-1(zzy009) |
| ZZY10435 | *C. nigoni*, Cni-shls-1(zzy010) IV, zzyIR10417 (IV, AF16>JU1421), zzyIs0734[Cbr-myo-2p::GFP; NeoR] IV | Cni-shls-1(zzy010) |
| ZZY10436 | *C. nigoni*, CNI13191(zzy011) IV, zzyIR10417 (IV, AF16>JU1421), zzyIs0734[Cbr-myo-2p::GFP; NeoR] IV | CNI13191(zzy011) |
| ZZY10437 | *C. nigoni*, CNI13191(zzy012) IV, zzyIR10417 (IV, AF16>JU1421), zzyIs0734[Cbr-myo-2p::GFP; NeoR] IV | CNI13191(zzy012) |
| ZZY0997 | *C. nigoni*, Cni-shls-1(zzy016) IV | Cni-shls-1(zzy016) |
| ZZY1008 | *C. briggsae*, she-1(v49) IV, Cbr-shls-1(zzy018) IV | Cbr-shls-1(zzy018) |
| ZZY10508 | *C. nigoni*, zzyDf018 IV, zzyIR10481 (IV, AF16>JU1421), zzyIs0734[Cbr-myo-2p::GFP; NeoR] IV | zzyDf018 |
| ZZY10509 | *C. nigoni*, zzyDf019 IV, zzyIR10481 (IV, AF16>JU1421), zzyIs0734[Cbr-myo-2p::GFP; NeoR] IV | zzyDf019 |
| ZZY10510 | *C. nigoni*, zzyDf020 IV, zzyIR10481 (IV, AF16>JU1421), zzyIs0734[Cbr-myo-2p::GFP; NeoR] IV | zzyDf020 |
| ZZY10511 | *C. nigoni*, zzyDf021 IV, zzyIR10481 (IV, AF16>JU1421), zzyIs0734[Cbr-myo-2p::GFP; NeoR] IV | zzyDf021 |
| ZZY10515 | *C. nigoni*, g15727(zzy029) IV, zzyIR10481 (IV, AF16>JU1421), zzyIs0734[Cbr-myo-2p::GFP; NeoR] IV | g15727(zzy029) |
| ZZY10516 | *C. nigoni*, g15727(zzy030) IV, zzyIR10481 (IV, AF16>JU1421), zzyIs0734[Cbr-myo-2p::GFP; NeoR] IV | g15727(zzy030) |
| ZZY10517 | *C. nigoni*, zzyDf022 IV, zzyIR10481 (IV, AF16>JU1421), zzyIs0734[Cbr-myo-2p::GFP; NeoR] IV | zzyDf022 |
| ZZY10518 | *C. nigoni*, zzyDf023 IV, zzyIR10481 (IV, AF16>JU1421), zzyIs0734[Cbr-myo-2p::GFP; NeoR] IV | zzyDf023 |
| ZZY10519 | *C. nigoni*, zzyDf024 IV, zzyIR10481 (IV, AF16>JU1421), zzyIs0734[Cbr-myo-2p::GFP; NeoR] IV | zzyDf024 |
| ZZY10520 | *C. nigoni*, zzyDf025 IV, zzyIR10481 (IV, AF16>JU1421), zzyIs0734[Cbr-myo-2p::GFP; NeoR] IV | zzyDf025 |
| ZZY10521 | *C. nigoni*, Cni-neib-1(zzy031) IV, zzyIR10481 (IV, AF16>JU1421), zzyIs0734[Cbr-myo-2p::GFP; NeoR] IV | Cni-neib-1(zzy031) |
| ZZY10522 | *C. nigoni*, Cni-neib-1(zzy032) IV, zzyIR10481 (IV, AF16>JU1421), zzyIs0734[Cbr-myo-2p::GFP; NeoR] IV | Cni-neib-1(zzy032) |
| ZZY1073 | *C. nigoni*, Cni-neib-1(zzy034) IV | Cni-neib-1(zzy034) |
| ZZY1095 | *C. nigoni*, Cni-neib-1(zzy035) IV | Cni-neib-1(zzy035) |
| ZZY1096 | *C. nigoni*, Cni-neib-1(zzy034), Cni-shls-1(zzy036) IV | Cni-neib-1(zzy034), Cni-shls-1(zzy036) |
| ZZY1097 | *C. nigoni*, Cni-neib-1(zzy035), Cni-shls-1(zzy037) IV | Cni-neib-1(zzy035), Cni-shls-1(zzy037) |
| ZZY1098 | *C. nigoni*, Cni-shls-1(zzy038) IV | Cni-shls-1(zzy038) |
| **Knock-in Strain** |  |  |
| ZZY1005 | *C. briggsae*, zzyIs1005[cel-eft-3p::Cni-shls-1::tbb-2 3'UTR; NeoR] IV | zzyIs1005 |
| ZZY1006 | *C. briggsae*, zzyIs1006[cel-eft-3p::Cni-shls-1::tbb-2 3'UTR; NeoR] IV | zzyIs1006 |
| ZZY1009 | *C. briggsae*, she-1(V49), zzyIs1006[cel-eft-3p::Cni-shls-1::tbb-2 3'UTR; NeoR] IV | zzyIs1005 |
| ZZY1010 | *C. briggsae*, she-1(V49), zzyIs1006[cel-eft-3p::Cni-shls-1::tbb-2 3'UTR; NeoR] IV | zzyIs1006 |
| ZZY1033 | *C. briggsae*, she-1(V49), zzyIs1033[Cbr-shls-1_motif_change_to_Cni-shls-1] IV | zzyIs1033 |
| ZZY1119 | *C. briggsae*, she-1(V49), zzyIs1119[mCherry::Cbr-shls-1] IV | zzyIs1119 |
| ZZY1122 | *C. nigoni*, zzyIs1122[gfp::Cni-shls-1] IV | zzyIs1122 |
| **Extrachromosomal Array Strain** |  |  |
| ZZY10439 | *C. nigoni*, zzyEx1001[cel-eft-3p::Cni-shls-1::tbb-2 3'UTR] | zzyEx1001 |
| ZZY10440 | *C. nigoni*, zzyEx1002[cel-eft-3p::Cni-shls-1::tbb-2 3'UTR] | zzyEx1002 |
| ZZY10443 | *C. briggsae*, she-1(V49), zzyEx1003[cel-eft-3p::Cni-shls-1::tbb-2 3'UTR] | zzyEx1003 |
| ZZY10444 | *C. briggsae*, she-1(V49), zzyEx1004[cel-eft-3p::Cni-shls-1::tbb-2 3'UTR] | zzyEx1004 |
| ZZY10445 | *C. briggsae*, *she-1*(V49), *zzyEx1005[cel-eft-3p::Cni-shls-1::tbb-2 3'UTR]* | *zzyEx1005* |
| ZZY10473 | *C. briggsae*, *she-1*(V49), *zzyEx1008[cel-eft-3p::Cbr-shls-1::tbb-2 3'UTR]* | *zzyEx1008* |
| ZZY10474 | *C. briggsae*, *she-1*(V49), *zzyEx1009[cel-eft-3p::Cbr-shls-1::tbb-2 3'UTR]* | *zzyEx1009* |
| ZZY10496 | *C. nigoni*, *zzyEx1016[cel-eft-3p::cbr-shls-1::tbb-2 3'UTR]* | *zzyEx1016* |
| ZZY10497 | *C. nigoni*, *zzyEx1017[cel-eft-3p::cbr-shls-1::tbb-2 3'UTR]* | *zzyEx1017* |
| ZZY10523 | *C. nigoni*, *zzyEx1016[g15728p::g15728]* | *zzyEx1016* |
| ZZY10524 | *C. nigoni*, *zzyEx1017[g15728p::g15728]* | *zzyEx1017* |
| ZZY10525 | *C. briggsae*, *she-1*(V49),<br>*zzyEx1018[g15728p::g15728]* | *zzyEx1018* |
| ZZY10526 | *C. briggsae*, *she-1*(V49),<br>*zzyEx1019[g15728p::g15728]* | *zzyEx1019* |
| ZZY10529 | *C. nigoni*, *zzyEx1016[g15728p::g15728]*, *Cni-neib-1(zzy034)*, *Cni-shls-1(zzy036) IV* | *zzyEx1016* |
| ZZY10530 | *C. nigoni*, *zzyEx1017[g15728p::g15728]*, *Cni-neib-1(zzy034)*, *Cni-shls-1(zzy036) IV* | *zzyEx1017* |
| ZZY10533 | *C. briggsae*, *she-1*(V49),<br>*zzyEx1018[g15728p::g15728]*, *zzyIs1006[cel-eft-3p::Cni-shls-1::tbb-2 3'UTR; NeoR] IV* | *zzyEx1018* |
| ZZY10534 | *C. briggsae*, *she-1*(V49),<br>*zzyEx1019[g15728p::g15728]*, *zzyIs1006[cel-eft-3p::Cni-shls-1::tbb-2 3'UTR; NeoR] IV* | *zzyEx1019* |

**Movie S1 (separate file). Rapid degradation of *Cbr-*SHLS-1 during the early embryogenesis in F1 hybrids from *C. briggsae* mothers (*mCherry::Cbr-shls-1(+/+)*) and *C. nigoni* fathers (*gfp::Cni-shls-1(+/+)*).**

The time (hh:mm:ss) for each frame and the scale bar are shown at the top-right and bottom-right corners, respectively.

**Movie S2 (separate file). *Cbr-*SHLS-1 expression during embryogenesis in *C. briggsae* with heterozygous *mCherry::Cbr-shls-1(+/-)*.**

*mCherry::Cbr-shls-1* allele was inherited from *C. briggsae* mothers homozygous for *mCherry::Cbr-shls-1(+/+)*. The time (hh:mm:ss) for each frame and the scale bar are shown at the top-right and bottom-right corners, respectively.

**Movie S3 (separate file). *Cni-*SHLS-1 expression during embryogenesis in *C. nigoni* with heterozygous *gfp::Cni-shls-1(+/-)*.**

*gfp::Cni-shls-1* allele was inherited from *C. nigoni* fathers homozygous for *gfp::Cni-shls-1(+/+)*. The time (hh:mm:ss) for each frame and the scale bar are shown at the top-right and bottom-right corners, respectively.

**Movie S4 (separate file). Absence of *Cbr-*SHLS-1 expression during embryogenesis in F1 hybrids from *C. briggsae* fathers (*mCherry::Cbr-shls-1(+/+)*) and *C. nigoni* (*gfp::Cni-shls-1(+/+)*).**

The time (hh:mm:ss) for each frame and the scale bar are shown at the top-right and bottom-right corners, respectively.

**Movie S5 (separate file). *Cbr-*SHLS-1 expression during embryogenesis in *C. briggsae* with heterozygous *mCherry::Cbr-shls-1(+/-)*.**

*mCherry::Cbr-shls-1* allele was inherited from *C. briggsae* fathers homozygous for mCherry::*Cbr-shls-1(+/+)*. The time (hh:mm:ss) for each frame and the scale bar are shown at the top-right and bottom-right corners, respectively.

**Movie S6 (separate file). *Cni-*SHLS-1 expression during embryogenesis in *C. nigoni* with heterozygous *gfp::Cni-shls-1(+/-)*.**

*gfp::Cni-shls-1* allele was inherited from *C. nigoni* mothers homozygous for *gfp::Cni-shls-1(+/+)*. The time (hh:mm:ss) for each frame and the scale bar are shown at the top-right and bottom-right corners, respectively.

**Movie S7 (separate file). No degradation of *Cbr-*SHLS-1 during embryogenesis in F1 hybrids from *C. briggsae* mothers (*mCherry::Cbr-shls-1(+/+)*) and *Cni-neib-1(-/-)* mutant fathers.**

The time (hh:mm:ss) for each frame and the scale bar are shown at the top-right and bottom-right corners, respectively.

**Movie S8 (separate file). No degradation of *Cbr-*SHLS-1 during embryogenesis in F1 hybrids from *C. briggsae* fathers (*mCherry::Cbr-shls-1(+/+)*) and *Cni-neib-1(-/-)* mutant mothers.**

The time (hh:mm:ss) for each frame and the scale bar are shown at the top-right and bottom-right corners, respectively.

